# Genome-wide identification of novel long non-coding RNAs and their possible roles in hypoxic zebrafish brain

**DOI:** 10.1101/2020.07.10.181842

**Authors:** Bodhisattwa Banerjee, Debaprasad Koner, David Karasik, Nirmalendu Saha

**Affiliations:** Musculoskeletal Genetics Laboratory, The Azrieli Faculty of Medicine, Bar-Ilan University, Safed 1311502, Israel; Biochemical Adaptation Laboratory, Department of Zoology, North-Eastern Hill University, Shillong 793022, India

**Keywords:** long non-coding RNAs, hypoxia, Alzheimer’s disease, brain regions, zebrafish

## Abstract

Long non-coding RNAs (lncRNAs) are the master regulators of numerous biological processes. Hypoxia causes oxidative stress with severe and detrimental effects on brain function and acts as a critical initiating factor in the pathogenesis of Alzheimer’s disease (AD). From the RNA-Seq in the forebrain (Fb), midbrain (Mb), and hindbrain (Hb) regions of hypoxic and normoxic zebrafish, we identified novel lncRNAs, whose potential *cis* targets showed involvement in neuronal development and differentiation pathways. Under hypoxia, several lncRNAs and mRNAs were differentially expressed. Co-expression studies indicated that the Fb and Hb regions’ potential lncRNA target genes were involved in the AD pathogenesis. In contrast, those in Mb (*cry1b, per1a, cipca*) were responsible for regulating circadian rhythm. We identified specific lncRNAs present in the syntenic regions between zebrafish and humans, possibly functionally conserved. We thus identified several conserved lncRNAs as the probable regulators of AD genes (*adrb3b, cav1, stat3, bace2, apoeb, psen1, s100b*).

## 1. Introduction

Non-coding RNAs (ncRNAs) were previously considered as “transcriptional noise” and deemed as “non-functional”. Recent technical advances in the high-throughput genomic platforms have revealed that despite active transcription of about 85% of the human genome [1,2], only 1 – 2% of it is protein-coding (PC). Thus, the vast majority of transcripts are represented by ncRNAs. Two major classes of ncRNAs have been classified according to their length: small ncRNAs (under 200 nt) and long non-coding RNAs (lncRNAs) (200 nt to as long as several kilobase pairs). Like the mRNAs, most of the lncRNAs undergo splicing, 5’ capping, 3’ polyadenylation and are usually transcribed by RNA polymerase II. Like coding transcripts, lncRNAs, too, have epigenetic markers [3] and may contain polymorphisms [4]. The lncRNAs are emerging as critical regulators of major biological processes impacting development, differentiation, and disease [5,6].

Hypoxia is the condition where there is an inadequate oxygen supply in tissues. Approximately 20% of the human body’s energy budget is consumed by the brain [7], and thus adequate supply of oxygen is critical for the normal functioning of the neurons. It has been suggested that hypoxia plays an essential role in the initiation of Alzheimer’s’ disease (AD) pathogenesis as hypoxic conditions increase free radical production in the electron transport chain in the mitochondria leading to increased oxidative stress [8,9].

The zebrafish is a small, hardy freshwater fish native to South Asian countries like India, Nepal, Pakistan, and Bangladesh. Originally zebrafish was used as a model organism for the study of vertebrate development. However, in recent decades large-scale genetic screens were able to identify hundreds of mutant phenotypes similar to human diseases, including the AD [8,10,11]. Several reported studies support the zebrafish model’s viability as an alternative way for a better understanding of neurodegenerative diseases, such as AD [12]. Nevertheless, for studies on all these diseases and AD, the focal spot is primarily on the PC genes. However, similar studies using the zebrafish to probe lncRNA function are still in infancy, mostly because of a lack of proper annotation and systematic survey.

To date, only a few studies have systematically determined the lncRNA landscape in the zebrafish. Ulitsky et al. [13] used chromatin marks and poly(A) site mapping and RNA-seq data from three developmental stages and reported more than 550 distinct lncRNAs in zebrafish. Among which 29 lncRNAs showed sequence similarity with their mammalian orthologs. Similarly, two more studies by Pauli et al. [14] and Kaushik et al. [15] identified 1133 and 442 long non-coding transcripts from eight developmental stages of zebrafish embryogenesis and different adult tissues, respectively.

However, there are several lacunae in these previous studies. Moreover, no data are available on the lncRNAs expressed in the fish brain’s sub-regions. Therefore, in this study, we used a strand-specific RNA-sequencing (ssRNA-Seq) method to identify and characterize novel lncRNAs (previously unreported) and their potential target PC genes in three different brain regions of the zebrafish. Furthermore, to elucidate the roles of these lncRNAs and their potential target genes under hypoxia, we constructed co-expression networks of lncRNAs and mRNAs in these brain sub-regions. There is very little sequence conservation of many lncRNAs between species, mostly conservation is restricted to short sequence stretches [16]. However, many lncRNAs were derived from syntenic loci between different species and thus thought to be functionally conserved [16]. Therefore, we ascertained the conserved syntenic loci of important novel lncRNAs between zebrafish and humans in this current study.

## 2. Results

### 2.1 Genome-wide identification of novel lncRNAs

From the adult zebrafish brain regions (forebrain, midbrain, and hindbrain) RNA-Seq library, a total of ~530 million reads were obtained. GC content and Q30 content in these data averaged at 43.8% and 93.94%, respectively (Supplementary Table S1). An average of 78.77%, 79.83%, and 81.03% of reads mapped uniquely to the zebrafish genome (GRCz11) in the Fb, Mb, and Hb regions, respectively (Supplementary Table S2). We have used stringent criteria to develop a pipeline for identifying novel lncRNAs (Fig. 1A). StringTie with --merge option was used to merge all the GTF files from the three brain sub-regions and produce a non-redundant set of transcripts of the zebrafish brain. Using the filtering pipeline (Fig. 1A), a total of 35060 lncRNA transcripts were identified. Furthermore, using the BLASTN search (E value 1e^−5^) against the lncRNA sequences of the ZFLNC database resulted in the detection of 4172 novel lncRNA transcripts (3527 novel lncRNA genes) (Supplementary file S1). The ZFLNC database is the most comprehensive database of zebrafish lncRNAs to date. The data consists of resources from NCBI, ENSEMBL, NONCODE, zflncRNApedia, and literature [17]. Thus, the transcripts without a BLAST hit against the ZFLNC data were considered as “novel”. Genomic locations of the novel lncRNA transcripts were identified using Ensembl genome browser (Fig. 1B). Among these novel lncRNAs majority (58%) was intergenic, 27% were intronic, and 12% were antisense (Fig. 1C). The majority (45%) of the novel lncRNAs identified in this study were 400 – 700 bp in length, whereas 16% of these were longer than 1000 bp in size (Fig. 1D). We identified the common lncRNAs in the Fb, Mb, and Hb regions and observed that 204 lncRNAs were mutual among all three regions (Supplementary fig. S1).

**Fig. 1.**
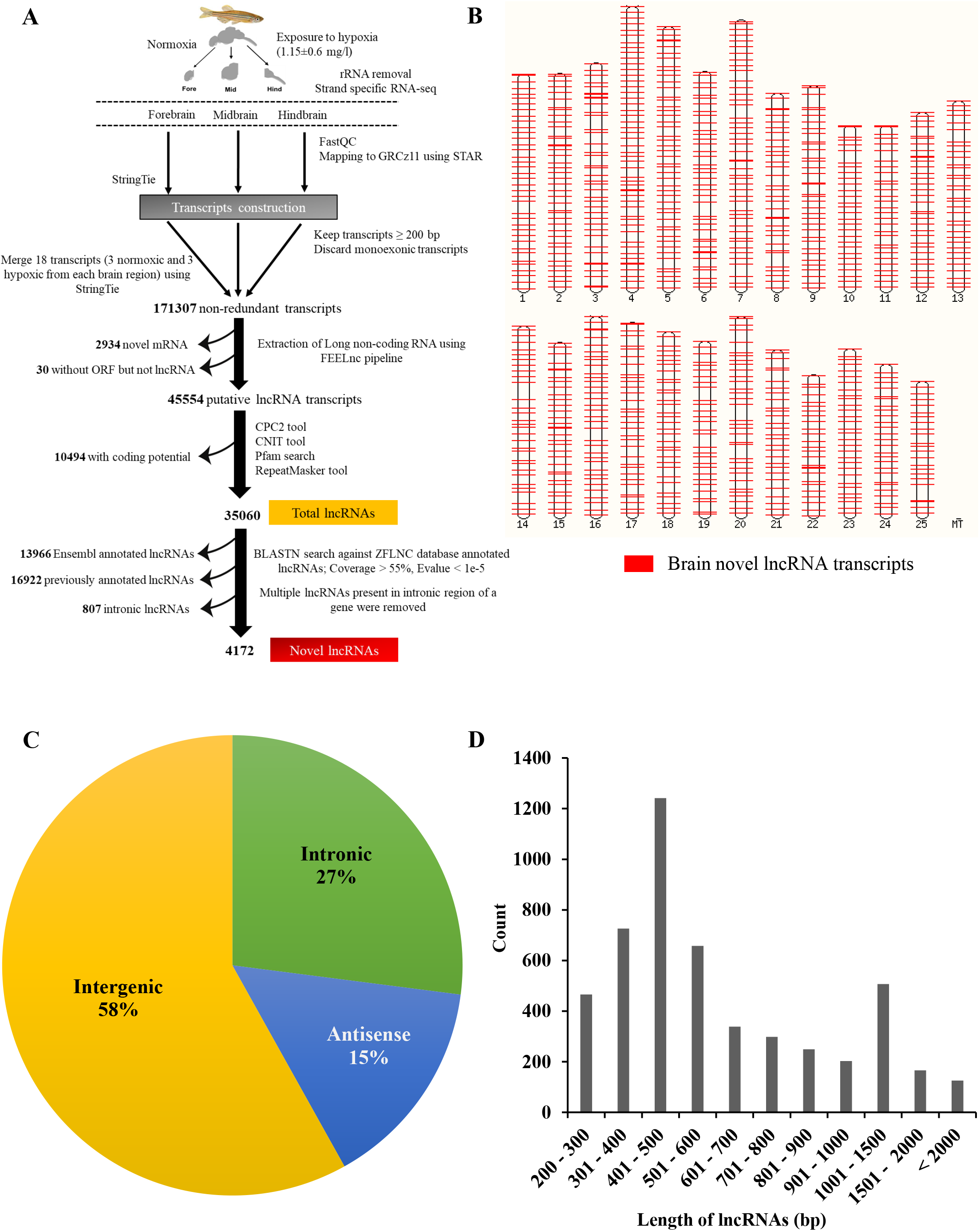
(A) Pipeline for identifying novel lncRNAs from the normoxic and hypoxic zebrafish brain regions. (B) Genomic distribution of the novel lncRNA transcripts. (C) Numbers of lncRNAs in each of the four main classes, as defined by their genomic location relative to neighboring or overlapping genes. (D) Length distribution of lncRNAs. X-axis: length of lncRNAs in bp; Y-axis: count.

### 2.2 Functional annotation of the novel lncRNAs

The PC genes present 100 kb up or downstream of the novel lncRNAs were considered *cis* target genes. We could identify 2564 potential *cis* target PC genes (Supplementary file S2). To obtain further insight into the functional role of these lncRNAs, we performed GO (Biological process) and KEGG pathway enrichment analysis (Fig. 2, Supplementary fig. 2 and Supplementary file S3 – S4). The top 20 terms enriched in the GO biological process include nervous system development, neurogenesis, multicellular organism development, system development. Similarly, three functional categories were enriched in the KEGG pathway analysis, including the calcium signaling pathway, apelin signaling pathway, and cell adhesion molecules (CAMs). We identified 158 lncRNA and mRNA pairs that co-expressed.

**Fig. 2.**
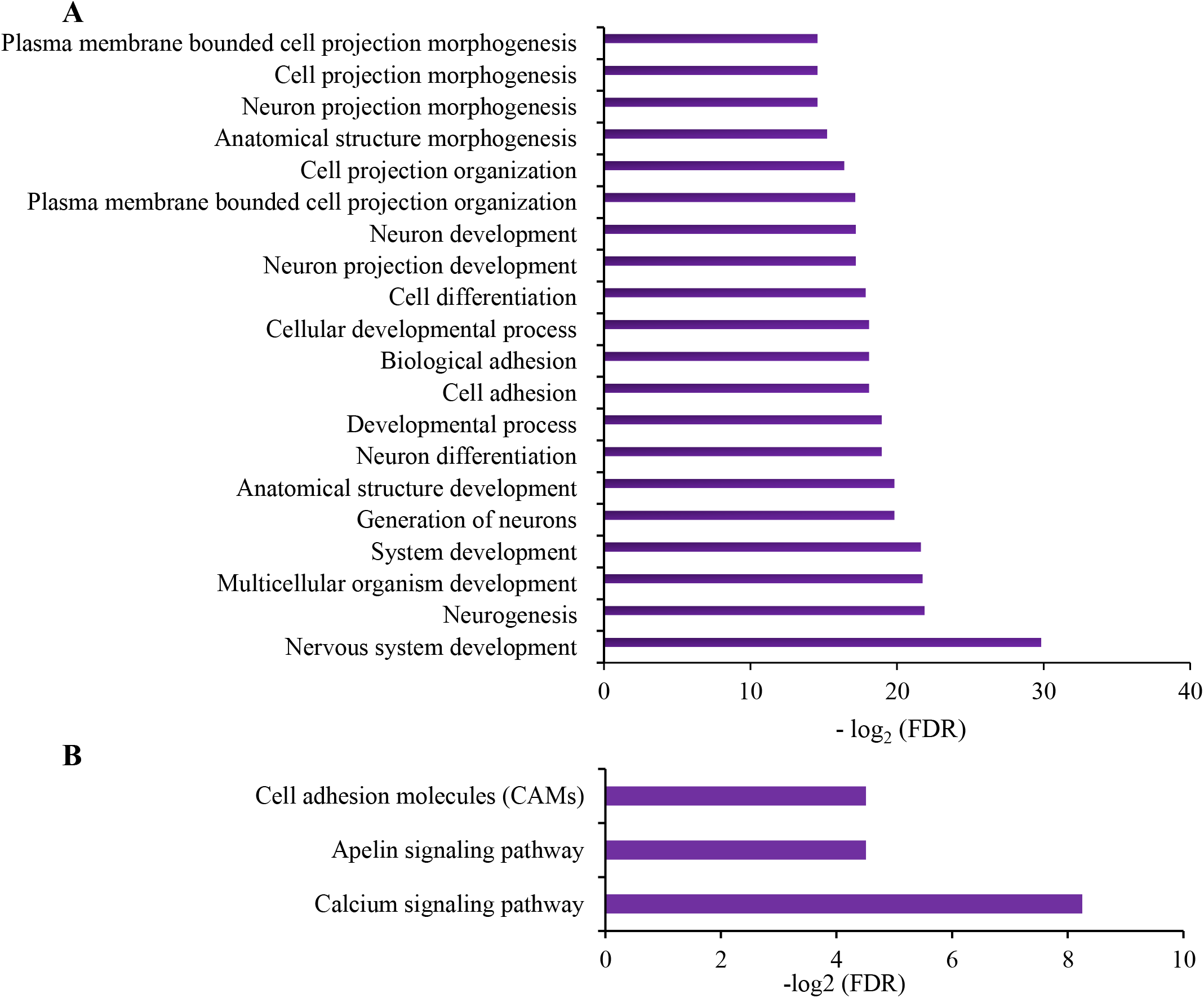
Enrichment analysis of the *cis* target PC genes using (A) GO biological process (B) KEGG pathway

### 2.3 Differential expression of novel lncRNAs and mRNAs under hypoxia

Differential expression of the lncRNA and mRNA genes in three brain sub-regions were measured using the edgeR tool. The lncRNAs and mRNAs in the hypoxic group (1.1±0.6 mg/l of dissolved oxygen) of fish were compared to the normoxic (7.6±0.4 mg/l of dissolved oxygen) group, and the genes with fold change >2 and FDR <0.05 were considered as differentially expressed (DE). A total of 836, 920, and 1647 novel lncRNA (Supplementary file S5A – S5C) and 2723, 2379, and 6267 PC and Ensembl annotated lncRNA DE genes were detected in the Fb, Mb, and Hb regions, respectively (Supplementary file S6A – S6C). Many DE PC genes were implicated in positive or negative regulation of neurodegenerative diseases like AD (i.e., *adrb3b, apoeb, psen1, s100b, bace2, stat3, cav1, apoa1a, hspb1*, etc.). Volcano plots of the DE (up- or downregulated) lncRNAs (Fig. 3A) and mRNAs (Fig. 3B) identified the genes with large fold changes that are also statistically significant. Furthermore, we compared the number of DE lncRNAs and mRNAs between the three brain sub-regions. Venn diagram showed that 470 DE novel lncRNAs (Fig. 3C) were shared between all the three brain regions, whereas 1209 DE mRNAs (Fig. 3D) were common in all three brain regions. We used hierarchical cluster analysis to illustrate the expression patterns of DE lncRNAs and DE mRNAs in each brain region. The heatmap of the top 100 DE lncRNA and mRNA genes showed a clear separation between the normoxic and hypoxic groups in the three brain sub-regions (Figs. 3E and 3F).

**Fig. 3.**
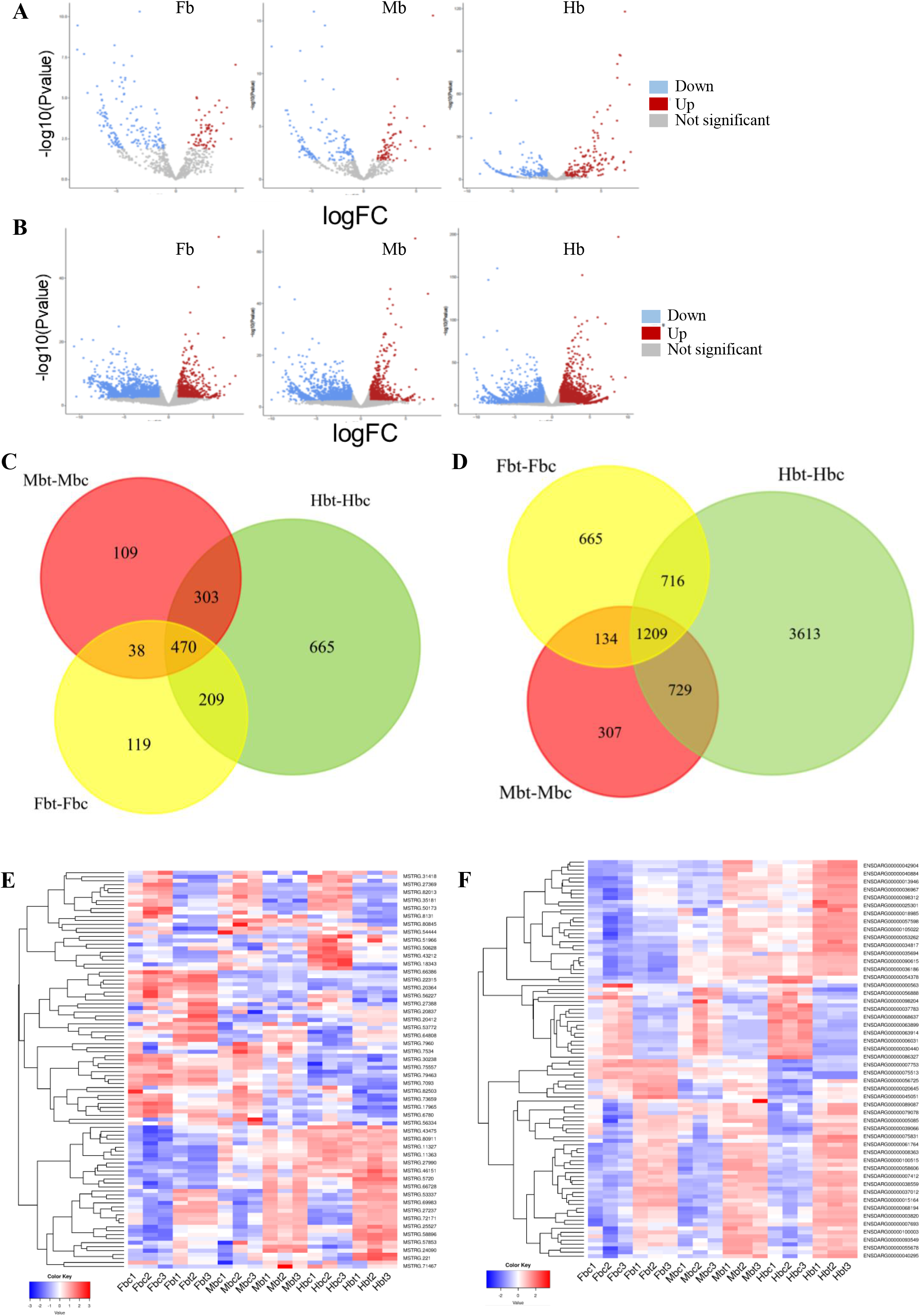
Volcano plot for identification statistically significant DE (up- or downregulated) (A) lncRNA genes and (B) mRNA genes. In this plot the genes are colored if they pass the thresholds for FDR and log fold change, red if they are upregulated and blue if they are downregulated. (C) Venn diagram showing the DE lncRNAs in three different brain regions between hypoxic and normoxic fish; (D) Venn diagram showing the DE mRNAs in three different brain regions between hypoxic and normoxic fish. Fbc: forebrain normoxia control, Fbt: forebrain hypoxia treated, Mbc: midbrain normoxia control, Mbt: midbrain hypoxia treated, Hbc: hindbrain normoxia control, Hbt: hindbrain hypoxia treated. Hierarchical clustering with heatmap of top 100 (E) DE lncRNA genes and (F) DE mRNA genes in all three brain regions between hypoxic and normoxic fish. The red color represents upregulated expression, and the blue color represents downregulated expression.

Moreover, enrichment analysis of the DE mRNAs in each brain sub-region was performed using the GO biological process, KEGG, and Disease.ZFIN databases. The GO enrichment analysis of the Fb region showed that the most significantly upregulated mRNAs were associated with developmental process, anion transport, multicellular organism development (Fig. 4A and Supplementary Table S3). Only one KEGG pathway, namely the MAPK signaling pathway, was enriched in the Fb region (Fig. 4B and Supplementary Table S4). In the Mb region, circadian regulation of gene expression, circadian rhythm, development process, and anatomical structure development were the four most significantly enriched GO terms (Fig. 4D and Supplementary Table S6). A single KEGG pathway, ECM-receptor interaction, was enriched in the Mb region (Fig. 4E and Supplementary Table S7). At the same time, 15 GO terms were enriched in the Hb region. Eight were significantly upregulated, including organic anion transport, organic acid transport, and chondrocyte morphogenesis (Fig. 4G and Supplementary Table S9). Four KEGG pathways were also enriched in the Hb region (Fig. 4H and Supplementary Table S10). The top DE mRNAs in all the three regions when using Disease.ZFIN were significantly enriched in type 2 diabetes mellitus, myocardial infarction (Figs. 4C, 4F, 4I, and Supplementary Table S5, S8, and S11). Additionally, in both Fb and Hb regions, DE mRNAs were significantly enriched for AD (Fig. 4C and 4I).

**Fig. 4.**
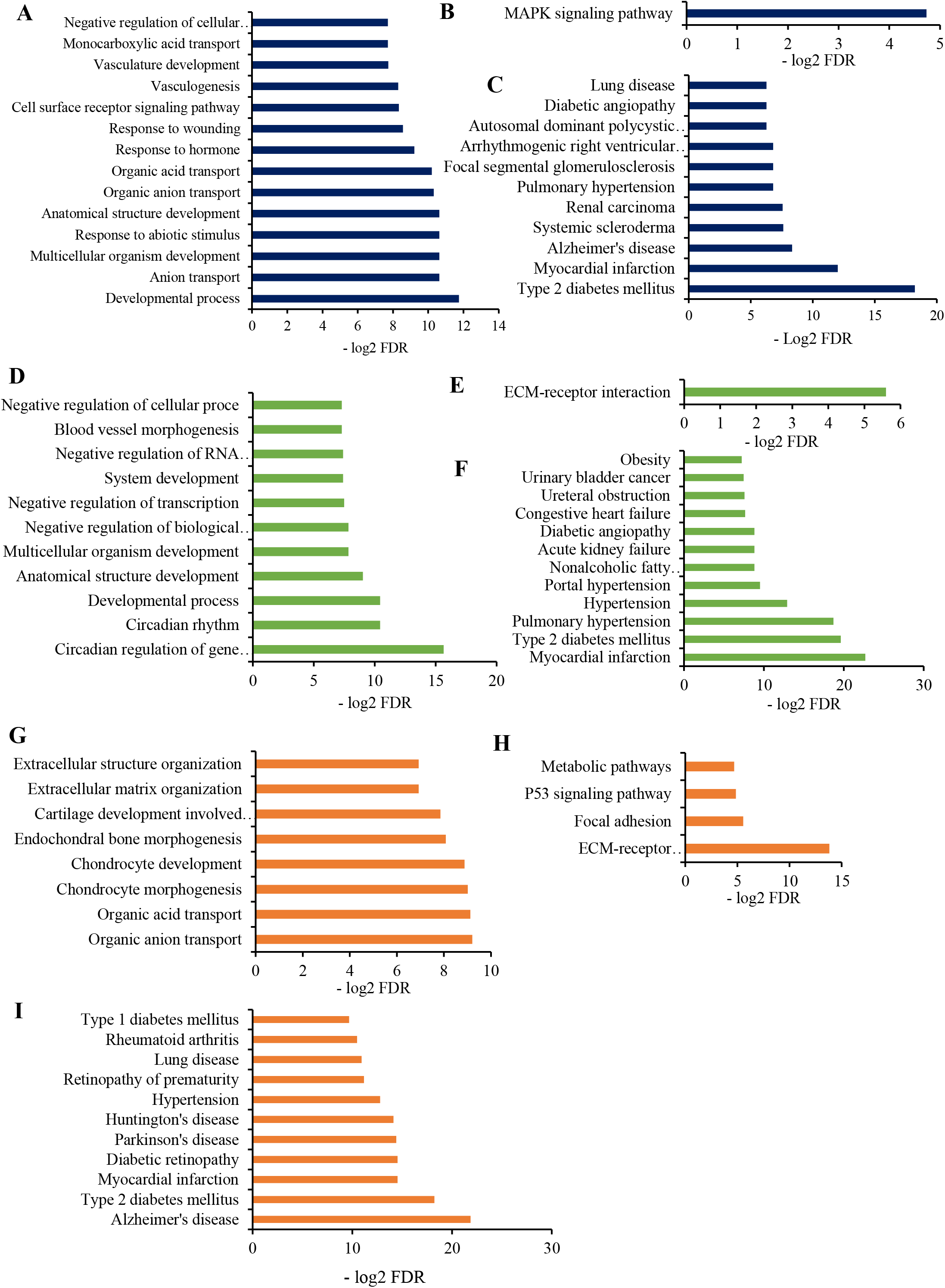
Enrichment analysis of the DE PC genes in Fb region (A) using GO biological process (B) using KEGG pathway (C) using Disease.ZFIN, in Mb region (D) using GO biological process (E) using KEGG pathway (F) using Disease.ZFIN, in Hb region (G) using GO biological process (H) using KEGG pathway (I) using Disease.ZFIN.

### 2.4 Construction of co-expression network of lncRNAs and mRNAs

Using weighted correlation network analysis in WGCNA (18 samples), we created clusters (modules) of highly correlated lncRNA and PC genes in each brain region. A uniquely colored label was assigned to each of the modules (Fig. 5A, 5B, 5C). In the Fb region, the lncRNAs and the correlated mRNAs were grouped into 9 clusters, with a range of 26 (magenta) to 720 (turquoise) genes in each cluster module (Fig. 5A). All the modules produced a co-expression network between the DE novel lncRNAs and mRNAs. However, the turquoise module had the highest number of such interactions. Based on the turquoise module, 170 lncRNAs produced a co-expression network with 560 mRNAs. Enrichment analysis of the turquoise module against the Disease.ZFIN database showed that genes were enriched for eight diseases, including Alzheimer’s disease, type 2 diabetes mellitus, myocardial infarction, and obesity (Supplementary fig. S1A and Supplementary Table S12).

**Fig. 5.**
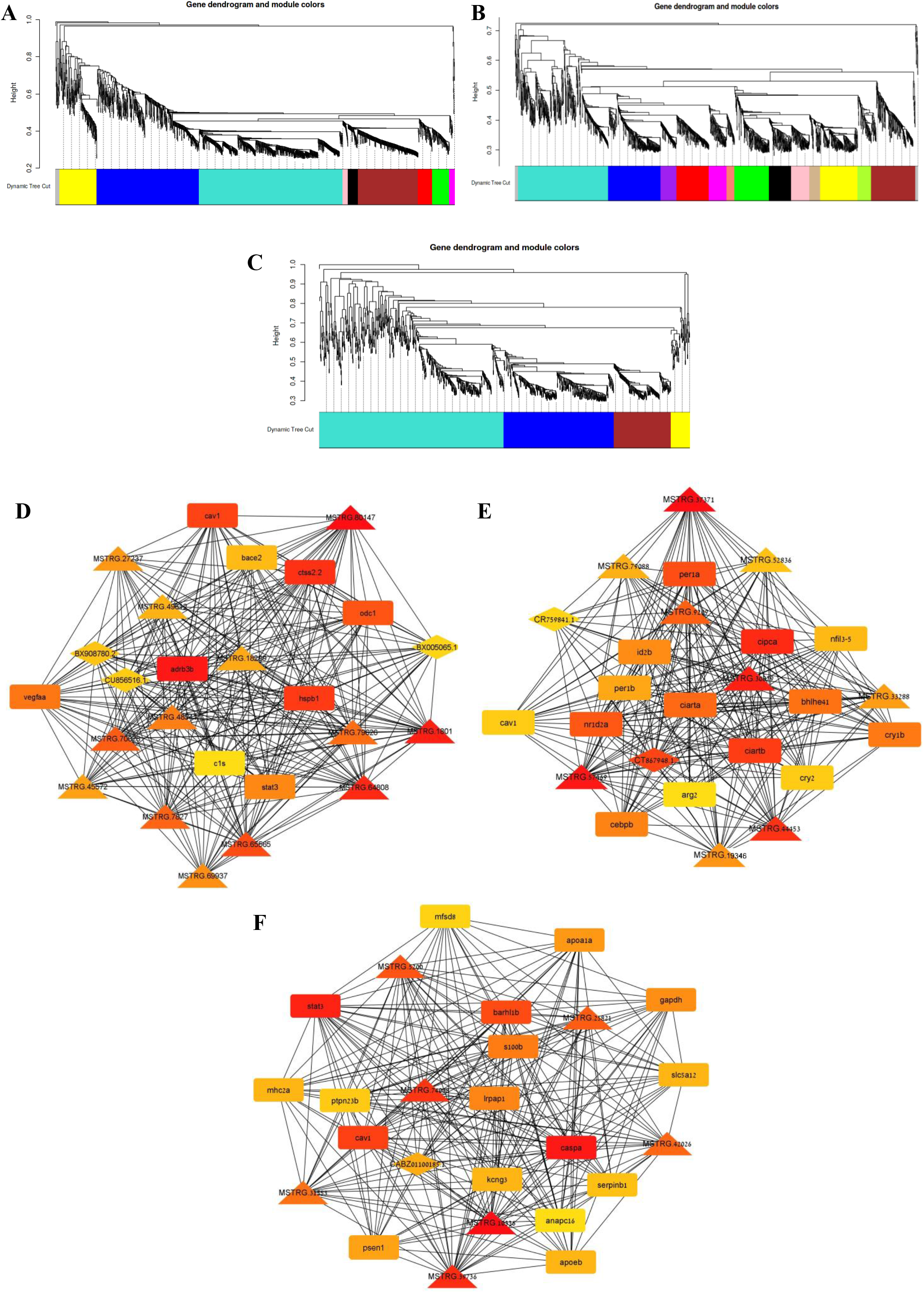
Different modules generated by the WGCNA. Hierarchical clustering dendrogram of co-expressed lncRNAs and mRNAs in modules. The colored strips below each dendrogram demonstrate the module designation identified through the clusters of co-expressed genes. (A) Fb region (B) Mb region (C) Hb region. The 25 top scoring hub gene co-expression network based on the MCC algorithm of lncRNA and PC gene. A redder color represents the higher-scoring ones. (D) Fb region (E) Mb region, (F) Hb region. The triangles denote the novel lncRNAs; the diamonds denote annotated lncRNAs; the squares denote PC genes.

Similarly, the DE lncRNAs and mRNAs were grouped into 13 and 4 modules, and the turquoise and blue modules were used for the generation of co-expression network in the Mb and Hb regions, respectively. Remarkably, the DE co-expression network genes in all three brain regions were enriched for the myocardial infarction, type 2 diabetes mellitus when we used Disease.ZFIN database (Supplementary figures S1A – S1C and Supplementary Tables S12 – S14). In addition to it, the genes in the co-expression networks in the Hb region were significantly enriched for AD (Supplementary fig. S1C and Supplementary Tables S14).

### 2.5 Potential regulatory lncRNA subnetwork

The network centrality was applied in this study to identify potential regulatory lncRNAs in the co-expression network. A Cytoscape plugin, cytoHubba, was used to determine the critical nodes and the hub genes in all the three brain regions’ co-expression networks. cytoHubba uses eleven analysis methods to compute the node scores, among which MCC has a better performance. The top 25 hub genes based on the MCC algorithm were selected for each brain region. The hub gene network in the Fb region consisted of 13 novel lncRNAs and 9 PC genes, and 3 annotated lncRNA (Fig. 5D). Majority of the PC genes (*adrb3b, ctss2.2, hspb1, cav1, vegfaa*, *stat3*, *bace2*) in the hub were involved in AD pathogenesis. The lncRNAs MSTRG.80147, MSTRG.1801, MSTRG.64808, MSTRG.65565, MSTRG.79626, MSTRG.48541, MSTRG.69937, and MSTRG.27237 had top scores and thus were likely to be the key regulators of these genes.

Similarly, 9 novel lncRNA, 14 PC and two annotated lncRNA genes were present in the Mb region’s hub genes subnetwork (Fig. 5E). Six lncRNA genes, MSTRG.37371, MSTRG.30829, MSTRG.57469, MSTRG.44453, MSTRG.19346, MSTRG.9209 had the highest score thus most likely acted as the potential regulators of the PC genes. Most of the PC genes (*cipca*, *ciartb*, *ciarta*, *per1a*, *nr1d2a*, *cry1b*, *per1b*, *cry2*) were involved in the circadian rhythm and circadian regulation of gene expression.

Moreover, in the Hb region, seven novel lncRNAs (i.e., MSTRG.10336, MSTRG.74008, MSTRG.39736, MSTRG.5200, MSTRG.25821, MSTRG.42026, MSTRG.31553) and 17 PC genes and a single annotated lncRNA (CABZ01100185.1) created the hub gene network (Fig. 5F). The PC genes were mostly implicated in AD, and among these eight PC genes, i.e., *stat3, caspa, cav1, s100b, lrpap1, apoa1a, apoeb, psen1* had the highest score.

### 2.6 Syntenic conservation and genomic features of the hub lncRNAs

Comparative genomic analysis between zebrafish and humans showed multiple potential regulatory lncRNAs present in the hub gene network appear in syntenic positions without any or little sequence conservation. The syntenic regions were determined using the Ensembl genome browser. Among the brain novel lncRNAs, 394 were located in syntenic regions (Supplementary Table S16). We had also identified the genomic features of the critical lncRNAs (synteny conserved with humans) using the histone H3 lysine 4 trimethylation (H3K4me3), histone H3 lysine 36 trimethylation (H3K36me3), CAGE-seq, and JASPAR TFBS (transcription factor binding sites) tracks in the UCSC genome browser. In the Fb region, four potential regulatory lncRNAs, MSTRG.64808, MSTRG.65565, MSTRG.48541, MSTRG.27237 were present in the syntenic areas with human chromosomes (Figs. 6A – 6H). Among these, MSTRG.64808 and MSTRG.48541 even have sequence conservation with human chromosome 5 and chromosome 8, respectively (Figs. 6B and 6F). Similarly, in the Mb region, two lncRNAs, MSTRG.30829 and MSTRG.19346, shared a syntenic region with humans (Figs. 7A – 7D). Moreover, four lncRNAs MSTRG.42026, MSTRG.25821, MSTRG.31553, and MSTRG.5200 present in the Hb region’s hub gene network also had synteny with human chromosomes (Figs. 7E – 7L). The genomic features of the potentially conserved lncRNA genes are shown in Supplementary Table S17. The majority of the lncRNAs were intergenic.

**Fig. 6.**
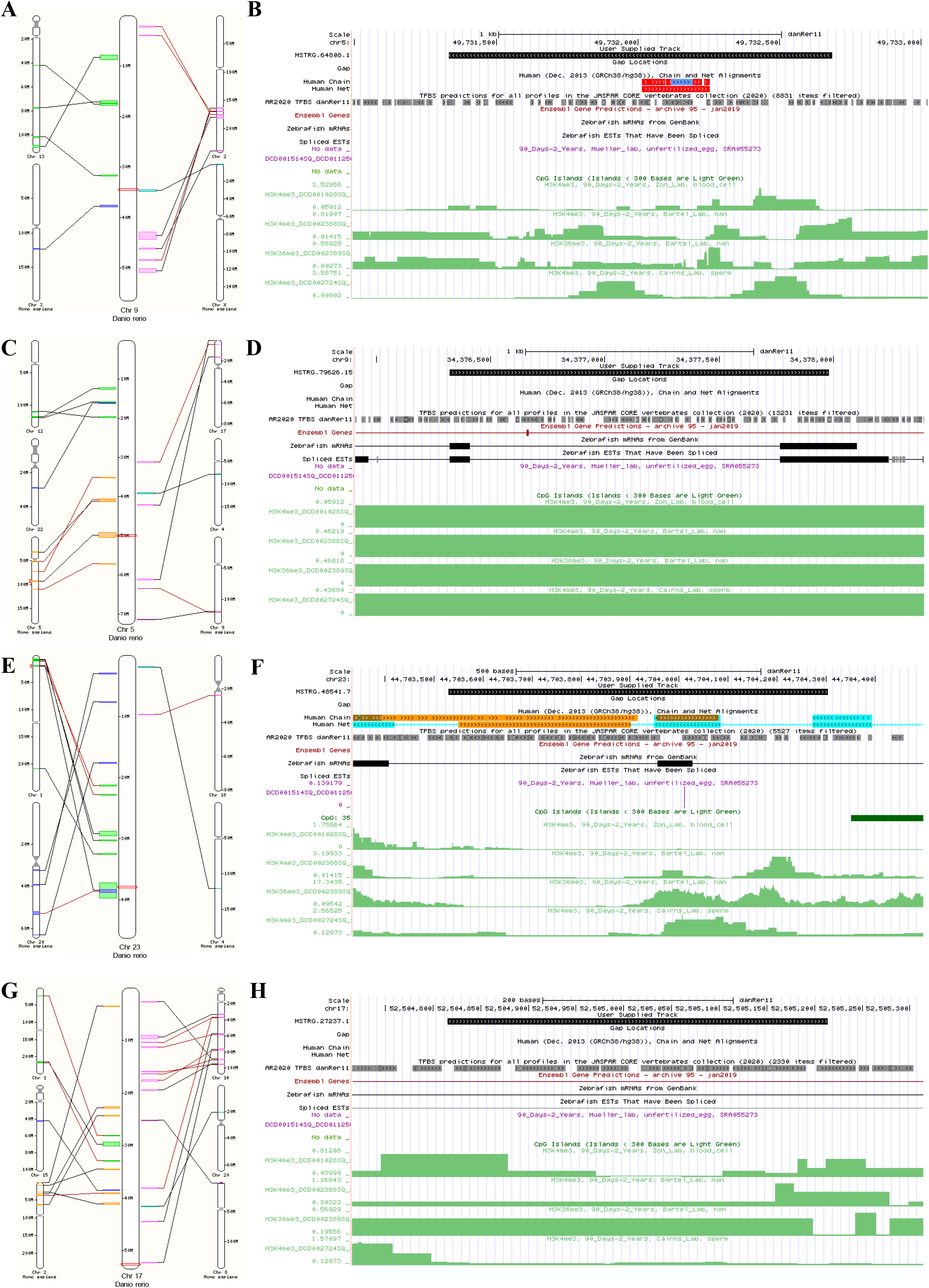
Synteny and genomic features of novel lncRNAs in Fb region (A), (C), (E), (G), The center chromosome represents the zebrafish chromosome, and the smaller chromosomes show syntenic regions with humans. Blocks are colored according to the chromosome number on the second species. These blocks are connected by lines. The small red box marks the location of the gene of interest and its homologue. (B), (D), (F), (H) Genomic features of the novel lncRNAs from the UCSC genome browser.

**Fig. 7.**
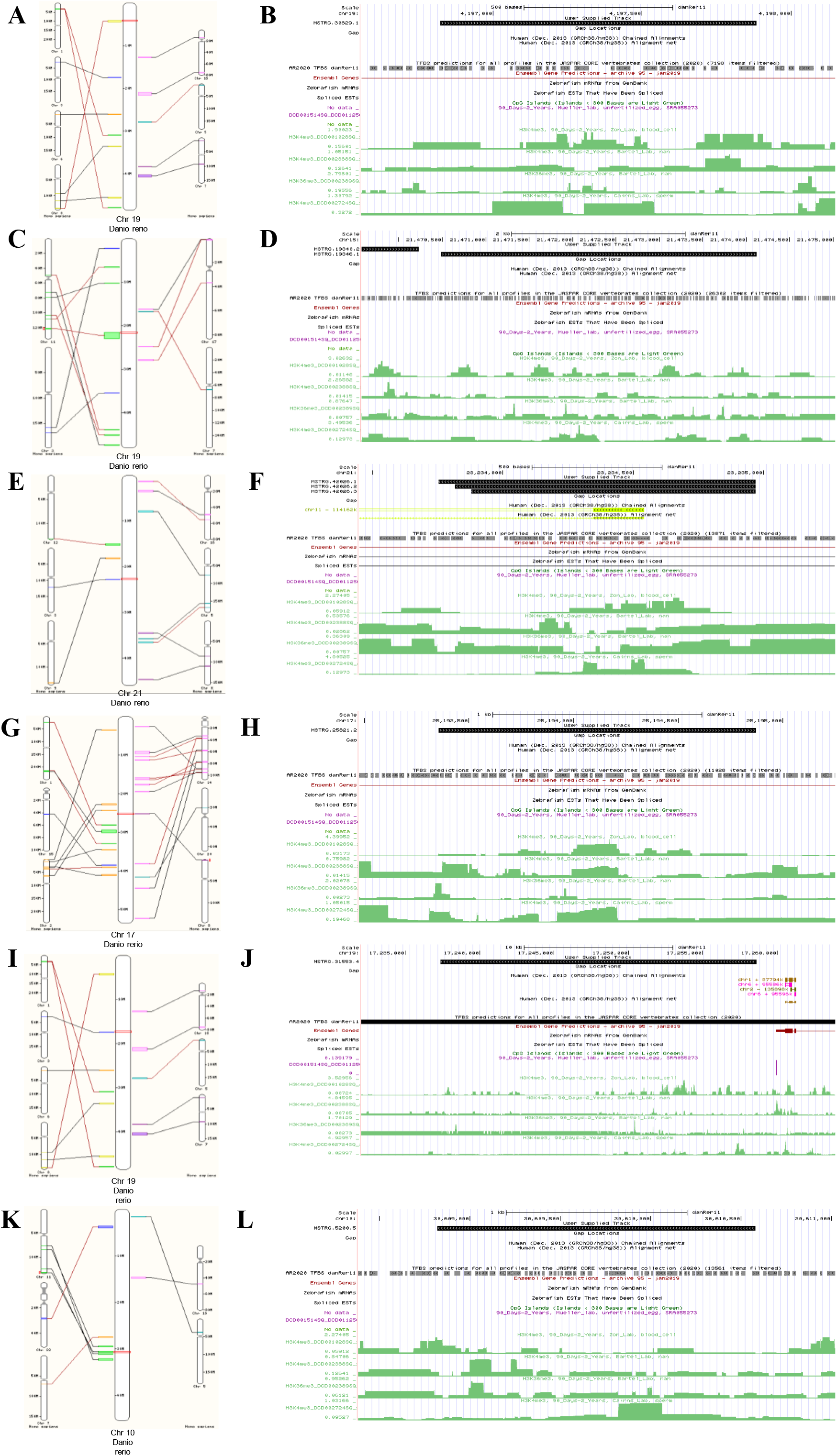
Synteny and genomic features of novel lncRNAs in Mb region (A), (C), The center chromosome represents the zebrafish chromosome, and the smaller chromosomes show syntenic regions with humans. Blocks are colored according to the chromosome number on the second species. These blocks are connected by lines. The small red box marks the location of the gene of interest and its homologue. (B), (D), Genomic features of the novel lncRNAs from the UCSC genome browser. Synteny and genomic features of novel lncRNAs in Hb region (E), (G), (I), (K), The center chromosome represents the zebrafish chromosome, and the smaller chromosomes show syntenic regions with humans. Blocks are colored according to the chromosome number on the second species. These blocks are connected by lines. The small red box marks the location of the gene of interest and its homologue. (F), (H), (J), (L) Genomic features of the novel lncRNAs from the UCSC genome browser.

### 2.7 Validation of RNA-Seq data by RT-qPCR

An equal number (four) of DE lncRNAs and DE mRNAs were randomly selected from the Fb, Mb, and Hb regions, and their expression was validated by RT-qPCR analysis. The expression patterns of the selected lncRNAs and mRNAs showed a similar trend in these two methods. These results (Fig. 8) validate the reliability of the RNA-Seq data.

**Fig. 8.**
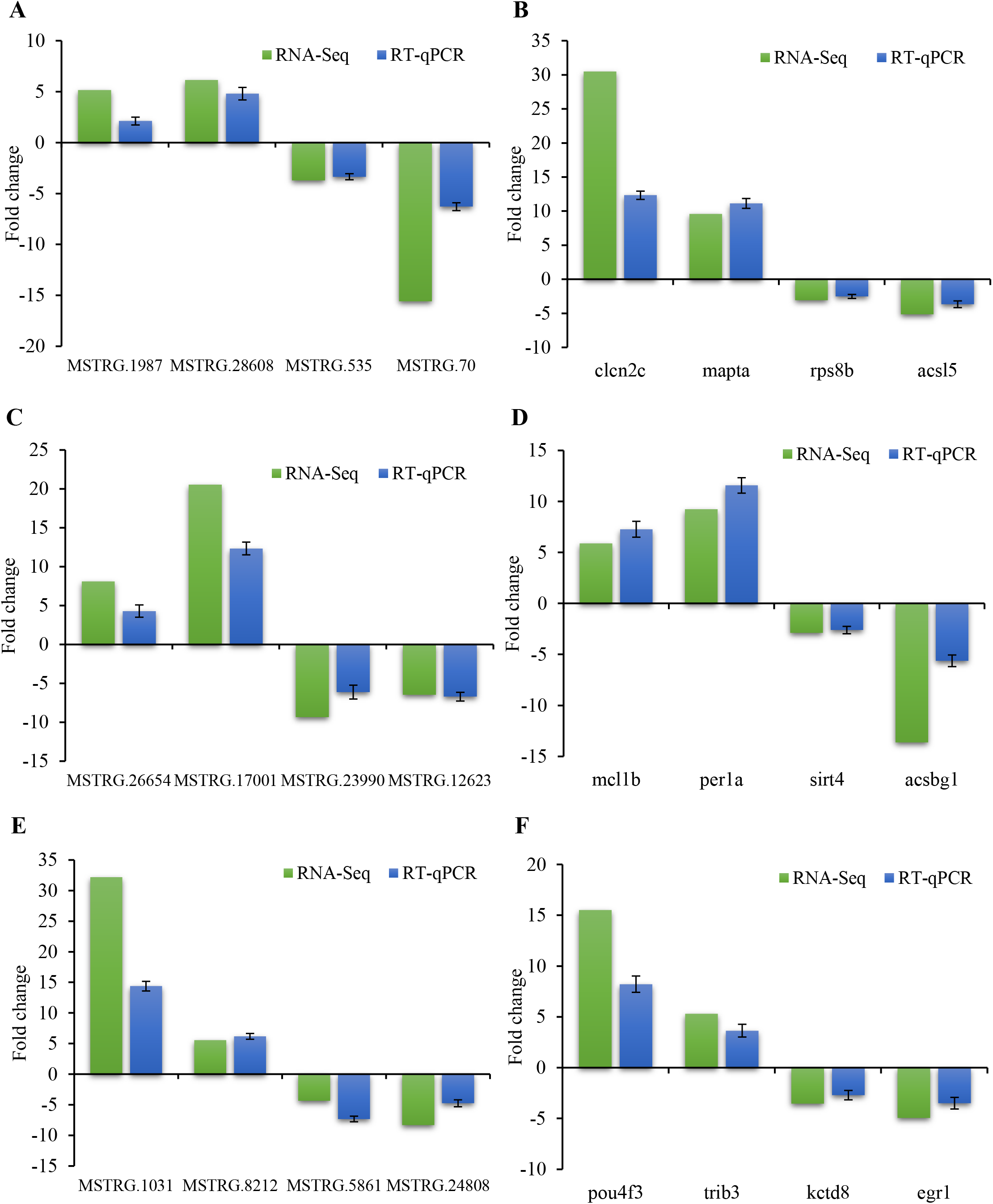
The RT-qPCR validation results of DE lncRNAs for (A) Fb, (C) Mb, and (E) Hb and DE mRNAs for (B) Fb, (D) Mb, and (F) Hb regions of the zebrafish brain, respectively.

## 3. Discussion

Among the small animal models, zebrafish has gained popularity as one of the valuable ones relevant for a study of vertebrate gene function in general. Thus, a high number (~70%) of human genes have at least a single zebrafish orthologue [18]. Moreover, zebrafish and the human brain have a high conservation in the neuroanatomical structures [19,20] and neurochemical pathways [21]. The zebrafish forebrain medial, dorsal, and lateral pallium has strong similarities with the humans’ amygdala, isocortex, and hippocampus, respectively [22,23]. Moreover, despite the size difference of particular zebrafish brain structures, major brain components, such as the diencephalon or the cerebellum, are highly conserved with the human brain [24]. Despite lacking the complex advanced cognitive behaviors evident in rodent models, zebrafish has proven to be very informative for studying human diseases, including AD [8]. Multiple studies have established the zebrafish as an important model for human disease-related PC genes. In contrast, knowledge of biology, function, and potential of lncRNAs as biomarkers and treatment targets in genetic diseases is still in its infancy. Very few studies have determined the lncRNA landscape in the zebrafish, and there are no reports on the sub-region specific expression of lncRNAs in the zebrafish brain. The only research elucidating brain lncRNAs of zebrafish was conducted by Kaushik et al. [15]. However, there were two significant problems in their study: (1) the method used for RNA isolation. They used the poly-A enrichment method. At times (but not always), lncRNAs are processed similarly to the mRNAs (e.g., capped, spliced, and polyadenylated) [25]. Hence, the RNA sample’s poly-A enrichment is not ideal for capturing all the lncRNAs expressed. (2) Also, they have used an older zebrafish genome version (Zv9) for alignment. The latest genome version and annotations should be used for accurate identification of lncRNAs [26]. So, to avoid these potential inaccuracies, in this study, we used ssRNA-Seq, the latest zebrafish genome (GRCz11), and a stringent filtering pipeline to systematically identify novel lncRNAs, and determined their differential expression pattern in three sub-regions of the brain under hypoxia. The fish model of brain hypoxia was obtained by the method described by Nik et al. [27].

A total of 2564 PC genes were identified as the potential *cis* targets of the novel lncRNAs in the total brain. GO Enrichment analysis of these targets showed that most genes were implicated in neuronal development, neurogenesis, multicellular organism development, and system development. KEGG pathway enrichment showed that most genes were enriched for calcium signaling and apelin signaling pathways. Numerous studies had revealed that multiple lncRNAs were involved in mammalian neurogenesis [28,29] as well as in neural development [30]. Neural physiology and viability are dependent upon the intracellular Ca^2+^-signaling. Disruptions in Ca^2+^ homeostasis lead to various neurodegenerative diseases, including AD [31]. The interplay between amyloid-β (Aβ) and Ca^2+^ is crucial in AD pathogenesis [31]. Binding of an important neuropeptide, apelin, to its receptor APJ triggers a diversity of physiological effects such as vasoconstriction and dilation, the control of blood pressure and blood flow, and had recently been implicated in neuroprotection [32]. Multiple studies have shown that apelin can prevent the production of Aβ and reduce its amounts by increasing its degradation [32]. From this study, it could be inferred that the novel lncRNA targets were mostly implicated in neural functioning and AD pathogenesis.

Consequently, from this study, it could be inferred that most of the novel lncRNAs detected in this study were involved in the regulatory role of neurodevelopmental procedures. We identified a largely region-specific expression of the lncRNAs. Altogether, the expression profile of the Fb, Mb, and Hb region was distinct. Although a relatively small number of novel lncRNAs (204) were shared among all three different regions indicating that these lncRNAs might have critical regulatory roles for maintenance of all these regions, this was consistent with a significant number of studies showing that lncRNAs have brain region-specific expression pattern [15,33–35]. The knowledge that the functional roles of many lncRNAs in the brain are spatially specific is relevant to understanding their target PC genes’ functions.

Several studies had implicated brain hypoxia for the progression of AD. Many cardiovascular risk factors and vascular and traumatic brain injury have a strong correlation with AD pathogenesis [30]. Hypoxia in the human brain due to vascular function failure leads to a regionalized effect, extending progressively with decreased vascular functions [36]. The DE mRNA genes in the Fb and Mb regions in this study were also enriched for vasculogenesis, vasculature development, blood vessel morphogenesis, and diseases like myocardial infarction, pulmonary hypertension, and congestive heart failure, indicating a role in cardiovascular diseases. Most lncRNA expressions in the brain of AD patients show a decreasing pattern [37,38]. Our data follow a similar trend, where the majority of the novel lncRNAs were downregulated in hypoxic fish brains.

The co-expression analysis of the DE novel lncRNA, PC, and annotated lncRNA genes produced huge co-expression networks in the three brain regions. We identified the hub genes, or the potentially critical lncRNA and PC genes in a network. A single lncRNA had interactions with multiple PC genes as well as with other lncRNA genes. In the Fb region, the correlated DE mRNA targets showed a regulatory role in AD. The top 25 hub genes included several lncRNA genes, among which some of the top-scored lncRNAs, MSTRG.64808, MSTRG.48541, MSTRG.79626, MSTRG.1801, and MSTRG.27237 have correlated with the zebrafish orthologues (*adrb3b, cav1, ctss2.2*, and *stat3*) of the human *ADRB3* (adrenoceptor beta 3), *CAV1* (caveolin 1), *CTSS* (cathepsin S), and *STAT3* (signal transducer and activator of transcription 3), and all the mRNAs were significantly upregulated. The *ADRB3* gene product, a beta-3-adrenergic receptor, is predominantly located in adipose tissues and functions in thermogenesis regulation, oxygen consumption, and adipocyte lipolysis [39]. Nalls et al. [40] identified this gene as associated with the progression of late-onset AD. CAV1 interacts with the beta-site amyloid precursor protein cleaving enzyme 1 (BACE1) and protein kinase C (PKC) to modulate the β and γ secretase activity of amyloid precursor protein, respectively [41,42]. Like our study, increased *CAV1* gene expression has been reported in the AD patients’ brains [41]. Cerebral hypoxia causes neuroinflammation, which has also been linked to AD pathogenesis [43]. Consequently, various pro-inflammatory transcription factor genes such as *STAT3* and *CEBPB* (CCAAT enhancer binding protein beta) were activated in the AD and associated with the late-onset AD pathogenesis [44]. It is further emphasized by Reichenbach and colleagues [45], where they have demonstrated activation of STAT3 in both the AD mouse model and human AD patients. CEBPB is crucial in regulating cytokines and genes involved in the immune and inflammatory response. Interestingly, in our current study, *cebpb* was predicted as a target gene of two lncRNAs, MSTRG.30829 and MSTRG.19346, in the Mb region. The lncRNAs were significantly downregulated, whereas the potential target gene *cebpb* was upregulated multifold. These data altogether re-emphasized the importance of inflammatory responses in hypoxia-induced AD. Moreover, the *bace2* gene was also identified as the target of lncRNAs MSTRG.64808 and MSTRG.27237. The gene *bace2* is the zebrafish homolog of the human *BACE2* involved in AD pathogenesis [46]. Bace2 efficiently cleaves the amyloid beta A4 precursor protein (APP) in the middle of the Aβ domain [46]. Enrichment analysis of the co-network module using Disease.ZFIN database predicted AD as the top significantly enriched disease.

In the present study, the Hb region has a higher number of correlated DE lncRNA and mRNA genes than the other two regions due to its much higher DE genes. Furthermore, in the Hb region, the lncRNAs MSTRG.42026, MSTRG.25821, MSTRG.39736 were identified as the potential regulators of the *apoeb* gene. The gene *apoeb* is a zebrafish orthologue of human APOE (apolipoprotein E), one of the most substantial genetic risk factors for AD [47,48]. Although minimal research has yet been conducted on APOE functions in zebrafish, some studies elucidating the *apoea* [49] and *apoeb* [50] reported their expression in the retina, yolk syncytial layer, and microglial cell [51]. Additionally, MSTRG.5200, MSTRG.25821, MSTRG.74008, and MSTRG.10336 lncRNAs were in a co-expression network with multiple DE genes, namely *psen1* (presenilin 1), *s100b* (S100 calcium-binding protein, beta (neural)), and *caspa* (caspase a). Nik et al. [52] have shown an upregulation of *psen1, psen2, appa, appb*, and *bace2* in the zebrafish adult brain under hypoxia. Importantly, human brains showed a similar pattern of expression [53–55]. S100B acts as a pro-inflammatory cytokine, and several studies reported high levels of S100B in AD patients [56] and mouse models of AD [56]. Similarly, *s100b* expression in the zebrafish brain region was elevated. The zebrafish *caspa* is the orthologue of the human *CASP1* (caspase 1) gene, and its expression was several-fold upregulated in the hypoxic fish brain (Hb region) compared to their normoxic counterparts. Several studies had associated *CASP1* upregulation with activation of the inflammasome complex in mammals, which leads to AD pathogenesis [57–59].

Of note, in the Fb and Hb brain regions, expression of two essential genes present in the hub gene network, *hspb1*, and *apoa1a*, were significantly downregulated in hypoxic fish. The gene *hspb1* in the Fb shared the hub co-expression network with MSTRG.64808, MSTRG.48541, MSTRG. 18299, and MSTRG.1801. Furthermore, the gene *apoa1a* was potentially regulated by MSTRG.5200, MSTRG.25821, and MSTRG.74008 lncRNA genes. The gene products of the mammalian counterparts of these two genes (HSPB1 and APOA1) function as neuroprotectors. The HSPB1 inhibits the formation of the microtubule-associated protein tau fibril in AD by interacting with it [60]. Similarly, the HDL-cholesterol transporter APOA1 binding to the Aβ1-40 maintains the Aβ in solution, thus preventing its deposit in the brain and fibril formation [61]. The AD patients also have significantly lower APOA1 levels compared to the controls [62].

Curiously, in the Mb region, four lncRNAs, namely MSTRG.37371, MSTRG.19346, MSTRG.30829, and MSTRG.44453 were identified as the probable regulator of the circadian clock genes, *per1a, per1b, cry1b*, and *cry2* genes. The period (*PER1*, *PER2*, and *PER3*) and cryptochrome (*CRY1, CRY2*) genes coordinate biological rhythms by repressing the expression of *CLOCK* and *BMAL1* genes in a periodic manner [63]. Moreover, multiple studies have demonstrated that acute and chronic hypoxia changes the expression of *per1a* and *per1b* in zebrafish, whereas the hypoxia response is, in turn, regulated by the clock genes [63–65]. Neurodegenerative diseases, such as AD, affect the proper functioning of circadian clock genes and may disrupt the timely organization amid different areas of the brain [66].

The sequence-function relationship in classical non-coding RNAs was greatly influenced by comparative sequence analysis [67]. Identifying lncRNA evolution might discover essential regions in lncRNAs and highlight the features that drive their functions. However, the lncRNAs are poorly conserved across species [16]. However, their sequence conservation is less than PC genes, more than introns or random intergenic regions. A comparison of mammalian and zebrafish lncRNAs by Ulitsky et al. [16] showed that only 29 lincRNAs were conserved across species. However, they demonstrated that many lincRNAs in zebrafish were derived from syntenic loci between these species and might have similar functions. In the current study, we have used synteny annotations between zebrafish and humans from the Ensembl. Synteny is described as the conserved order of aligned genomic blocks between species, and it is calculated from the pairwise genome alignments. We identified 394 lncRNA transcripts (8.4%) from the total brain novel lncRNAs present in the syntenic loci with humans. Furthermore, we identified four, two, and four lncRNA genes in Fb, Mb, and Hb regions, respectively, from the hub gene co-expression networks located in the syntenic loci with humans. These lncRNAs were involved in the correlation network with several important DE PC genes implicated in AD pathogenesis. Therefore, these lncRNAs might be functionally conserved across species, despite their lack of sequence similarity.

## 4. Conclusion

In summary, we performed ssRNA-Seq in the three brain regions, namely Fb, Mb, and Hb of normoxic and hypoxic zebrafish and identified novel lncRNAs in the total brain as well as specific sub-regions. The functional enrichment studies of the potential *cis* targets of the novel lncRNAs suggested that they were involved in neuronal development and differentiation pathways. These genes were also identified as enriched for calcium and apelin signaling pathways. Under hypoxia, multiple lncRNAs and mRNAs were differentially expressed. The lncRNA expression pattern showed a difference between the brain regions. It could be inferred from the current study that in the Fb and Hb regions, the potential lncRNA targets were the genes implicated in the AD pathogenesis. In the Mb region, the potential target genes were involved in the circadian rhythm and regulation of circadian gene expression. From the present data, we identified multiple lncRNAs present in the syntenic genomic regions between zebrafish and humans, and thus potentially functionally conserved. We identified the lncRNAs MSTRG.64808, MSTRG.79626, MSTRG.48541, and MSTRG.27237 in the Fb region and lncRNAs MSTRG.25821, MSTRG.31553, MSTRG.5200, and MSTRG.42026 in the Hb region as the potential regulators of several AD genes (*adrb3b, cav1, ctss2.2, stat3, bace2, apoeb, psen1, s100b*, and *caspa*). However, further studies are necessary to elaborate on the specific functions of these lncRNAs in the pathogenesis of hypoxia-induced AD.

## 5. Materials and methods

### 5.1 Zebrafish maintenance and experimental procedures

The adult wild-type zebrafish were maintained in our laboratory as per the approved guidelines of the Institutional Animal Ethics Committee of North-Eastern Hill University. Zebrafish were bred and maintained at 28°C on a 14 h light/10 h dark cycle. 1-year-old adult zebrafish (AB strain) were used for all experiments (n =24). No sex differentiation of the fish was performed. The experiments conducted under low oxygen conditions were performed following the method described by Nik et al., 2014 [27]. Briefly, oxygen was depleted by bubbling nitrogen gas through the water, and dissolved oxygen level (DOL) was measured by a portable dissolved oxygen meter (Horiba Scientific, Japan). The DOL in the hypoxia group was 1.15±0.6 mg/l. Another group of fish (n = 12) was kept under normal oxygen conditions (6.6 ± 0.45 mg/l). The hypoxic conditions were maintained for 3 hours, and immediately after the treatment, three biological replicates, each consisting of 3 zebrafish, were used for isolation of RNA. A total of 18 fish (9 hypoxic and 9 normoxic) were sacrificed after anesthetizing with 0.03% tricaine (Sigma-Aldrich, Germany), and brains were separated and placed in ice-cold HBSS (1X Hank’s basal salt solution, 2.5 mM HEPES-KOH, pH 7.4, 35 mM glucose)(Sigma-Aldrich, USA). Three sub-regions of the brain were dissected, and RNA was isolated using the NucleoSpin RNA XS kit (Macherey-Nagel, Germany) following the manufacture’s protocol. The RNA quality and concentration were measured in the QIAxcel Advanced system (Qiagen, Germany) using QIAxcel RNA QC Kit v2.0 (Qiagen) and Qubit 3.0 fluorometer (Thermo Fisher Scientific, USA). The RNA samples with an RNA integrity score (RIS) value of more than eight were used in further processing.

### 5.2 Library construction and sequencing

Ribosomal RNA was depleted from the total RNA using the Ribo-Zero Gold rRNA removal kit (Human/Mouse/Rat) (Illumina, USA). The ribo-depleted RNA was used to create an ssRNA-Seq library, according to the TruSeq^®^ Stranded Total RNA sample preparation guide. The sequencing was performed on the NovaSeq 6000 system (Illumina) using 2×100 bp chemistry by Macrogen Inc., South Korea. The sequencing data have been deposited in the NCBI’s Gene Expression Omnibus with the GEO accession number **GSE153687**.

### 5.3 Transcriptome assembly and novel lncRNA detection

Reads with a quality score of ≥30 were selected for the downstream analysis. The raw reads were aligned to the latest zebrafish genome (GRCz11) from Ensembl using STAR v2.7.3 [69], and transcripts were assembled by StringTie [70] using the reference annotation from Ensembl. The transcripts of < 200 bp length and monoexonic were excluded from further analysis. Several stringent filtration steps were applied to obtain the lncRNAs (Fig. 1A). At first, the combined data from Stringtie --merge were used in the FEELnc tool [71] to get a potential list of lncRNAs. Next, the coding potential of the assigned lncRNAs was again evaluated using CPC2 [72] and CNIT [73] tool. Any transcript assigned as coding by any of the software were discarded. Furthermore, these lncRNAs were evaluated by Pfam search, and transcripts with E value ≥ 1e^−3^ were omitted from further analysis. Finally, the RepeatMasker tool was utilized to identify the sequences with repetitive regions, and those transcripts were excluded. The total lncRNAs obtained were matched against the lncRNAs from the ZFLNC database [17] using BLASTN (coverage ≥ 55% and E value ≤ 1e^−5^), and those without a match were deemed as candidate novel lncRNAs in zebrafish brain regions. Multiple lncRNAs located in intronic regions of a single gene were also discarded to produce final list of novel lncRNA transcripts. The novel lncRNAs were classified according to their localization and the direction of transcription of proximal RNA transcripts using the FEELnc_classifier module of the FEELnc pipeline [71].

### 5.4 Prediction of lncRNA targets

Potential target PC genes of the novel lncRNAs were predicted based on *cis* function. The closest PC gene 100 kb upstream and downstream of lncRNAs were screened using the FEELnc pipeline [71]. Functional enrichment of these target genes using the GO biological process and KEGG pathways were done using the ShinyGO v0.61 web-server [74].

### 5.5 Differential expression of lncRNAs

The raw read counts of the lncRNAs and mRNAs were determined by the featureCounts program [75], and differential expression analysis was performed using edgeR. Benjamini-Hochberg method for controlling the false discovery rate (FDR) was used for adjusting the *p-* value. Only the genes with FDR < 0.05 and fold change > 2 were considered differentially expressed (DE). Enrichment analysis of the DE lncRNA-targeted PC genes among GO biological process, KEGG pathways, and Disease.ZFIN databases were conducted using iDEP (integrated Differential Expression and Pathway analysis) web-server [76].

### 5.6 Co-expression network construction and functional enrichment analysis

The co-expression network of DE lncRNAs and mRNAs (18 samples) were constructed using Weighted Gene Co-expression Network Analysis (WGCNA) package [77] on the iDEP web-server. All parameters were set as the default. Different modules were detected using the dynamic tree cutting method, and the network of each module was visualized using Cytoscape v3.8.1 [78]. A Cytoscape plugin, cytoHubba [79] were used to select significant modules and analyze the network centrality, identifying the nodes perceived to be potential regulators of the underlying biological processes.

### 5.7 Validation of expression of lncRNAs

The differential expression profiles of lncRNAs and mRNAs between three brain regions were validated using a total of 12 lncRNAs and mRNAs (4 from each region) by qPCR analysis. The lncRNA and mRNA-specific primers (Supplementary Table S15) were designed by Primer-BLAST. Total RNA isolated earlier using the NucleoSpin RNA XS kit (Macherey-Nagel) was converted to the first-strand cDNA following the method described earlier [80]. qPCR reactions were performed in a StepOne plus Real time PCR system (Thermo Fisher Scientific) using SYBR^®^ Premix Ex Taq™ II (Takara Clontech, Japan). Melting curve analysis was used to re-confirm the amplification of only a single PCR product. Fold changes of genes in hypoxic fish compared to normoxic controls were calculated using the modified ΔΔCT method [81].

## Acknowledgments

This study was supported by a National Postdoctoral Fellowship project sanctioned to BB (PDF/2016/002013) by the SERB, New Delhi, India, and to NS by the National Agricultural Science Fund, Indian Council of Agricultural Research, New Delhi (ABA-7011/ 1018-19/240). NS acknowledges the FIST-II program (SR/FST/LS1-615/ 2014) by the DST, New Delhi. We thank Dr. Sumit Mukherjee, Bar-Ilan University, Israel, for his suggestions on bioinformatics works.

## References

[1] S. Djebali, C.A. Davis, A. Merkel, A. Dobin, T. Lassmann, A. Mortazavi, A. Tanzer, J. Lagarde, W. Lin, F. Schlesinger, C. Xue, G.K. Marinov, J. Khatun, B.A. Williams, C. Zaleski, J. Rozowsky, M. Röder, F. Kokocinski, R.F. Abdelhamid, T. Alioto, I. Antoshechkin, M.T. Baer, N.S. Bar, P. Batut, K. Bell, I. Bell, S. Chakrabortty, X. Chen, J. Chrast, J. Curado, T. Derrien, J. Drenkow, E. Dumais, J. Dumais, R. Duttagupta, E. Falconnet, M. Fastuca, K. Fejes-Toth, P. Ferreira, S. Foissac, M.J. Fullwood, H. Gao, D. Gonzalez, A. Gordon, H. Gunawardena, C. Howald, S. Jha, R. Johnson, P. Kapranov, B. King, C. Kingswood, O.J. Luo, E. Park, K. Persaud, J.B. Preall, P. Ribeca, B. Risk, D. Robyr, M. Sammeth, L. Schaffer, L.-H. See, A. Shahab, J. Skancke, A.M. Suzuki, H. Takahashi, H. Tilgner, D. Trout, N. Walters, H. Wang, J. Wrobel, Y. Yu, X. Ruan, Y. Hayashizaki, J. Harrow, M. Gerstein, T. Hubbard, A. Reymond, S.E. Antonarakis, G. Hannon, M.C. Giddings, Y. Ruan, B. Wold, P. Carninci, R. Guigó, T.R. Gingeras, Landscape of transcription in human cells, Nature. 489 (2012) 101–108. doi:10.1038/nature11233.

[2] M.J. Hangauer, I.W. Vaughn, M.T. McManus, Pervasive Transcription of the Human Genome Produces Thousands of Previously Unidentified Long Intergenic Noncoding RNAs, PLoS Genet. 9 (2013) e1003569. doi:10.1371/journal.pgen.1003569.

[3] S. Sati, S. Ghosh, V. Jain, V. Scaria, S. Sengupta, Genome-wide analysis reveals distinct patterns of epigenetic features in long non-coding RNA loci, Nucleic Acids Res. 40 (2012). doi:10.1093/nar/gks776.

[4] H. Ling, R. Spizzo, Y. Atlasi, M. Nicoloso, M. Shimizu, R.S. Redis, N. Nishida, R. Gafa, J. Song, Z. Guo, C. Ivan, E. Barbarotto, I. De Vries, X. Zhang, M. Ferracin, M. Churchman, J.F. van Galen, B.H. Beverloo, M. Shariati, F. Haderk, M.R. Estecio, G. Garcia-Manero, G.A. Patijn, D.C. Gotley, V. Bhardwaj, I. Shureiqi, S. Sen, A.S. Multani, J. Welsh, K. Yamamoto, I. Taniguchi, M.-A. Song, S. Gallinger, G. Casey, S.N. Thibodeau, L. Le Marchand, M. Tiirikainen, S.A. Mani, W. Zhang, R. V. Davuluri, K. Mimori, M. Mori, A.M. Sieuwerts, J.W.M. Martens, I. Tomlinson, M. Negrini, I. Berindan-Neagoe, J.A. Foekens, S.R. Hamilton, G. Lanza, S. Kopetz, R. Fodde, G.A. Calin, CCAT2, a novel noncoding RNA mapping to 8q24, underlies metastatic progression and chromosomal instability in colon cancer, Genome Res. 23 (2013) 1446–1461. doi:10.1101/gr.152942.112.

[5] R.A. Flynn, H.Y. Chang, Long Noncoding RNAs in Cell-Fate Programming and Reprogramming, Cell Stem Cell. 14 (2014) 752–761. doi:10.1016/j.stem.2014.05.014.

[6] J.J. Quinn, H.Y. Chang, Unique features of long non-coding RNA biogenesis and function, Nat. Rev. Genet. 17 (2016) 47–62. doi:10.1038/nrg.2015.10.

[7] S. Herculano-Houzel, Scaling of brain metabolism with a fixed energy budget per neuron: Implications for neuronal activity, plasticity and evolution, PLoS One. 6 (2011) e17514. doi:10.1371/journal.pone.0017514.

[8] M. Newman, E. Ebrahimie, M. Lardelli, Using the zebrafish model for Alzheimer’s disease research, Front. Genet. 5 (2014) 189. doi:10.3389/fgene.2014.00189.

[9] E.L. Bell, T.A. Klimova, J. Eisenbart, C.T. Moraes, M.P. Murphy, G.R.S. Budinger, N.S. Chandel, The Qo site of the mitochondrial complex III is required for the transduction of hypoxic signaling via reactive oxygen species production, J. Cell Biol. 177 (2007) 1029–1036. doi:10.1083/jcb.200609074.

[10] G.J. Lieschke, P.D. Currie, Animal models of human disease: Zebrafish swim into view, Nat. Rev. Genet. 8 (2007) 353–367. doi:10.1038/nrg2091.

[11] K. Dooley, Zebrafish: a model system for the study of human disease, Curr. Opin. Genet. Dev. 10 (2000) 252–256. doi:10.1016/S0959-437X(00)00074-5.

[12] E.M. Caramillo, D.J. Echevarria, Alzheimer’s disease in the zebrafish: Where can we take it?, Behav. Pharmacol. 28 (2017) 179–186. doi:10.1097/FBP.0000000000000284.

[13] I. Ulitsky, A. Shkumatava, C.H. Jan, H. Sive, D.P. Bartel, Conserved function of lincRNAs in vertebrate embryonic development despite rapid sequence evolution, Cell. 147 (2011) 1537–1550. doi:10.1016/j.cell.2011.11.055.

[14] A. Pauli, E. Valen, M.F. Lin, M. Garber, N.L. Vastenhouw, J.Z. Levin, L. Fan, A. Sandelin, J.L. Rinn, A. Regev, A.F. Schier, Systematic identification of long noncoding RNAs expressed during zebrafish embryogenesis., Genome Res. 22 (2012) 577–91. doi:10.1101/gr.133009.111.

[15] K. Kaushik, V.E. Leonard, S. Kv, M.K. Lalwani, S. Jalali, A. Patowary, A. Joshi, V. Scaria, S. Sivasubbu, Dynamic expression of long non-coding RNAs (lncRNAs) in adult zebrafish., PLoS One. 8 (2013) e83616. doi:10.1371/journal.pone.0083616.

[16] I. Ulitsky, A. Shkumatava, C.H. Jan, H. Sive, D.P. Bartel, Conserved Function of lincRNAs in Vertebrate Embryonic Development despite Rapid Sequence Evolution, Cell. 147 (2011) 1537–1550. doi:10.1016/j.cell.2011.11.055.

[17] X. Hu, W. Chen, J. Li, S. Huang, X. Xu, X. Zhang, S. Xiang, C. Liu, ZFLNC: a comprehensive and well-annotated database for zebrafish lncRNA, Database. 2018 (2018). doi:10.1093/database/bay114.

[18] K. Howe, M.D. Clark, C.F. Torroja, J. Torrance, C. Berthelot, M. Muffato, J.E. Collins, S. Humphray, K. McLaren, L. Matthews, S. McLaren, I. Sealy, M. Caccamo, C. Churcher, C. Scott, J.C. Barrett, R. Koch, G.-J. Rauch, S. White, W. Chow, B. Kilian, L.T. Quintais, J.A. Guerra-Assunção, Y. Zhou, Y. Gu, J. Yen, J.-H. Vogel, T. Eyre, S. Redmond, R. Banerjee, J. Chi, B. Fu, E. Langley, S.F. Maguire, G.K. Laird, D. Lloyd, E. Kenyon, S. Donaldson, H. Sehra, J. Almeida-King, J. Loveland, S. Trevanion, M. Jones, M. Quail, D. Willey, A. Hunt, J. Burton, S. Sims, K. McLay, B. Plumb, J. Davis, C. Clee, K. Oliver, R. Clark, C. Riddle, D. Elliott, G. Threadgold, G. Harden, D. Ware, S. Begum, B. Mortimore, G. Kerry, P. Heath, B. Phillimore, A. Tracey, N. Corby, M. Dunn, C. Johnson, J. Wood, S. Clark, S. Pelan, G. Griffiths, M. Smith, R. Glithero, P. Howden, N. Barker, C. Lloyd, C. Stevens, J. Harley, K. Holt, G. Panagiotidis, J. Lovell, H. Beasley, C. Henderson, D. Gordon, K. Auger, D. Wright, J. Collins, C. Raisen, L. Dyer, K. Leung, L. Robertson, K. Ambridge, D. Leongamornlert, S. McGuire, R. Gilderthorp, C. Griffiths, D. Manthravadi, S. Nichol, G. Barker, S. Whitehead, M. Kay, J. Brown, C. Murnane, E. Gray, M. Humphries, N. Sycamore, D. Barker, D. Saunders, J. Wallis, A. Babbage, S. Hammond, M. Mashreghi-Mohammadi, L. Barr, S. Martin, P. Wray, A. Ellington, N. Matthews, M. Ellwood, R. Woodmansey, G. Clark, J.D. Cooper, A. Tromans, D. Grafham, C. Skuce, R. Pandian, R. Andrews, E. Harrison, A. Kimberley, J. Garnett, N. Fosker, R. Hall, P. Garner, D. Kelly, C. Bird, S. Palmer, I. Gehring, A. Berger, C.M. Dooley, Z. Ersan-Ürün, C. Eser, H. Geiger, M. Geisler, L. Karotki, A. Kirn, J. Konantz, M. Konantz, M. Oberländer, S. Rudolph-Geiger, M. Teucke, C. Lanz, G. Raddatz, K. Osoegawa, B. Zhu, A. Rapp, S. Widaa, C. Langford, F. Yang, S.C. Schuster, N.P. Carter, J. Harrow, Z. Ning, J. Herrero, S.M.J. Searle, A. Enright, R. Geisler, R.H.A. Plasterk, C. Lee, M. Westerfield, P.J. de Jong, L.I. Zon, J.H. Postlethwait, C. Nüsslein-Volhard, T.J.P. Hubbard, H.R. Crollius, J. Rogers, D.L. Stemple, The zebrafish reference genome sequence and its relationship to the human genome, Nature. 496 (2013) 498–503. doi:10.1038/nature12111.

[19] T. Mueller, M.F. Wullimann, An Evolutionary Interpretation of Teleostean Forebrain Anatomy, Brain. Behav. Evol. 74 (2009) 30–42. doi:10.1159/000229011.

[20] M.F. Wullimann, B. Rupp, H. Reichert, M.F. Wullimann, B. Rupp, H. Reichert, The brain of the zebrafish Danio rerio: an overview, in: Neuroanat. Zebrafish Brain, Birkhäuser Basel, 1996: pp. 7–17. doi:10.1007/978-3-0348-8979-7_4.

[21] M.F. Wullimann, B. Rupp, H. Reichert, M.F. Wullimann, B. Rupp, H. Reichert, Introduction: neuroanatomy for a neurogenetic model system, in: Neuroanat. Zebrafish Brain, Birkhäuser Basel, 1996: pp. 1–5. doi:10.1007/978-3-0348-8979-7_1.

[22] M.R. Braford, Jr., Comparative Aspects of Forebrain Organization in the Ray-Finned Fishes: Touchstones or Not?, Brain. Behav. Evol. 46 (1995) 259–274. doi:10.1159/000113278.

[23] S. Saleem, R.R. Kannan, Zebrafish: an emerging real-time model system to study Alzheimer’s disease and neurospecific drug discovery, Cell Death Discov. 4 (2018) 45. doi:10.1038/s41420-018-0109-7.

[24] V. Tropepe, H.L. Sive, Can zebrafish be used as a model to study the neurodevelopmental causes of autism?, Genes, Brain Behav. 2 (2003) 268–281. doi:10.1034/j.1601-183X.2003.00038.x.

[25] M. Rutenberg-Schoenberg, A.N. Sexton, M.D. Simon, The Properties of Long Noncoding RNAs That Regulate Chromatin, Annu. Rev. Genomics Hum. Genet. 17 (2016) 69–94. doi:10.1146/annurev-genom-090314-024939.

[26] T. Weirick, G. Militello, R. Müller, D. John, S. Dimmeler, S. Uchida, The identification and characterization of novel transcripts from RNA-seq data., Brief. Bioinform. 17 (2015) 1–8. doi:10.1093/bib/bbv067.

[27] S.H.M. Nik, M. Newman, S. Ganesan, M. Chen, R. Martins, G. Verdile, M. Lardelli, Hypoxia alters expression of Zebrafish Microtubule-associated protein Tau (mapta, maptb) gene transcripts, BMC Res. Notes. 7 (2014) 1–9. doi:10.1186/1756-0500-7-767.

[28] D. Antoniou, A. Stergiopoulos, P.K. Politis, Recent advances in the involvement of long non-coding RNAs in neural stem cell biology and brain pathophysiology, Front. Physiol. 5 (2014) 155. doi:10.3389/fphys.2014.00155.

[29] A.D. Ramos, F.J. Attenello, D.A. Lim, Uncovering the roles of long noncoding RNAs in neural development and glioma progression, Neurosci. Lett. 625 (2016) 70–79. doi:10.1016/j.neulet.2015.12.025.

[30] T.C. Roberts, K. V. Morris, M.J.A. Wood, The role of long non-coding RNAs in neurodevelopment, brain function and neurological disease, Philos. Trans. R. Soc. B Biol. Sci. 369 (2014). doi:10.1098/rstb.2013.0507.

[31] A. Demuro, I. Parker, G.E. Stutzmann, Calcium signaling and amyloid toxicity in Alzheimer disease, J. Biol. Chem. 285 (2010) 12463–12468. doi:10.1074/jbc.R109.080895.

[32] J. Masoumi, M. Abbasloui, R. Parvan, D. Mohammadnejad, G. Pavon-Djavid, A. Barzegari, J. Abdolalizadeh, Apelin, a promising target for Alzheimer disease prevention and treatment, Neuropeptides. 70 (2018) 76–86. doi:10.1016/j.npep.2018.05.008.

[33] S.J. Liu, T.J. Nowakowski, A.A. Pollen, J.H. Lui, M.A. Horlbeck, F.J. Attenello, D. He, J.S. Weissman, A.R. Kriegstein, A.A. Diaz, D.A. Lim, Single-cell analysis of long non-coding RNAs in the developing human neocortex, Genome Biol. 17 (2016) 67. doi:10.1186/s13059-016-0932-1.

[34] M.N. Ziats, O.M. Rennert, Aberrant Expression of Long Noncoding RNAs in Autistic Brain, J. Mol. Neurosci. 49 (2013) 589–593. doi:10.1007/s12031-012-9880-8.

[35] B.M. Kadakkuzha, X.-A. Liu, J. McCrate, G. Shankar, V. Rizzo, A. Afinogenova, B. Young, M. Fallahi, A.C. Carvalloza, B. Raveendra, S. V. Puthanveettil, Transcriptome analyses of adult mouse brain reveal enrichment of lncRNAs in specific brain regions and neuronal populations., Front. Cell. Neurosci. 9 (2015) 63. doi:10.3389/fncel.2015.00063.

[36] J.L. Eberling, W.J. Jagust, B.R. Reed, M.G. Baker, Reduced temporal lobe blood flow in alzheimer’s disease, Neurobiol. Aging. 13 (1992) 483–491. doi:10.1016/0197-4580(92)90076-A.

[37] X. Zhou, J. Xu, Identification of Alzheimer’s disease–associated long noncoding RNAs, Neurobiol. Aging. 36 (2015) 2925–2931. doi:10.1016/j.neurobiolaging.2015.07.015.

[38] C. Shi, L. Zhang, C. Qin, Long non-coding RNAs in brain development, synaptic biology, and Alzheimer’s disease, Brain Res. Bull. 132 (2017) 160–169. doi:10.1016/j.brainresbull.2017.03.010.

[39] D.S. Meyers, S. Skwish, K.E.J. Dickinson, B. Kienzle, C.M. Arbeeny, β_3_-Adrenergic Receptor-Mediated Lipolysis and Oxygen Consumption in Brown Adipocytes from Cynomolgus Monkeys, J. Clin. Endocrinol. Metab. 82 (1997) 395–401. doi:10.1210/jcem.82.2.3738.

[40] M.A. Nalls, R.J. Guerreiro, J. Simon-Sanchez, J.T. Bras, B.J. Traynor, J.R. Gibbs, L. Launer, J. Hardy, A.B. Singleton, Extended tracts of homozygosity identify novel candidate genes associated with late-onset Alzheimer’s disease, Neurogenetics. 10 (2009) 183–190. doi:10.1007/s10048-009-0182-4.

[41] M.-J. Kang, Y.H. Chung, C.-I. Hwang, M. Murata, T. Fujimoto, I.-H. Mook-Jung, C.I. Cha, W.-Y. Park, Caveolin-1 upregulation in senescent neurons alters amyloid precursor protein processing, Exp. Mol. Med. 38 (2006) 126–133. doi:10.1038/emm.2006.16.

[42] C. Hattori, M. Asai, H. Onishi, N. Sasagawa, Y. Hashimoto, T.C. Saido, K. Maruyama, S. Mizutani, S. Ishiura, BACE1 interacts with lipid raft proteins, J. Neurosci. Res. 84 (2006) 912–917. doi:10.1002/jnr.20981.

[43] R. Lall, R. Mohammed, U. Ojha, What are the links between hypoxia and Alzheimer’s disease?, Neuropsychiatr. Dis. Treat. Volume 15 (2019) 1343–1354. doi:10.2147/NDT.S203103.

[44] X. Li, J. Long, T. He, R. Belshaw, J. Scott, Integrated genomic approaches identify major pathways and upstream regulators in late onset Alzheimer’s disease, Sci. Rep. 5 (2015) 12393. doi:10.1038/srep12393.

[45] N. Reichenbach, A. Delekate, M. Plescher, F. Schmitt, S. Krauss, N. Blank, A. Halle, G.C. Petzold, Inhibition of Stat3-mediated astrogliosis ameliorates pathology in an Alzheimer’s disease model, EMBO Mol. Med. 11 (2019). doi:10.15252/emmm.201809665.

[46] J. Walter, Fishing for function - distinct roles of Bace1 and Bace2 in Zebrafish development, J. Neurochem. 127 (2013) 435–437. doi:10.1111/jnc.12200.

[47] T.-P. V. Huynh, A.A. Davis, J.D. Ulrich, D.M. Holtzman, Apolipoprotein E and Alzheimer’s disease: the influence of apolipoprotein E on amyloid-β and other amyloidogenic proteins, J. Lipid Res. 58 (2017) 824–836. doi:10.1194/jlr.R075481.

[48] M.E. Belloy, V. Napolioni, M.D. Greicius, A Quarter Century of APOE and Alzheimer’s Disease: Progress to Date and the Path Forward, Neuron. 101 (2019) 820–838. doi:10.1016/j.neuron.2019.01.056.

[49] P.A. Raymond, L.K. Barthel, R.L. Bernardos, J.J. Perkowski, Molecular characterization of retinal stem cells and their niches in adult zebrafish, BMC Dev. Biol. 6 (2006) 36. doi:10.1186/1471-213X-6-36.

[50] Z. Pujic, Y. Omori, M. Tsujikawa, B. Thisse, C. Thisse, J. Malicki, Reverse genetic analysis of neurogenesis in the zebrafish retina, Dev. Biol. 293 (2006) 330–347. doi:10.1016/j.ydbio.2005.12.056.

[51] K.N. Veth, J.R. Willer, R.F. Collery, M.P. Gray, G.B. Willer, D.S. Wagner, M.C. Mullins, A.J. Udvadia, R.S. Smith, S.W.M. John, R.G. Gregg, B.A. Link, Mutations in Zebrafish lrp2 Result in Adult-Onset Ocular Pathogenesis That Models Myopia and Other Risk Factors for Glaucoma, PLoS Genet. 7 (2011) e1001310. doi:10.1371/journal.pgen.1001310.

[52] S.H. Moussavi Nik, L. Wilson, M. Newman, K. Croft, T.A. Mori, I. Musgrave, M. Lardelli, The BACE1-PSEN-AβPP regulatory axis has an ancient role in response to low oxygen/oxidative stress, J. Alzheimer’s Dis. 28 (2012) 515–530. doi:10.3233/JAD-2011-110533.

[53] R. Pluta, Pathological Opening of the Blood-Brain Barrier to Horseradish Peroxidase and Amyloid Precursor Protein following Ischemia-Reperfusion Brain Injury, Chemotherapy. 51 (2005) 223–226. doi:10.1159/000086924.

[54] R. De Gasperi, M. Sosa, S. Dracheva, G.A. Elder, Presenilin-1 regulates induction of hypoxia inducible factor-1α: altered activation by a mutation associated with familial Alzheimer’s disease, Mol. Neurodegener. 5 (2010) 38. doi:10.1186/1750-1326-5-38.

[55] X. Zhang, W. Le, Pathological role of hypoxia in Alzheimer’s disease, Exp. Neurol. 223 (2010) 299–303. doi:10.1016/j.expneurol.2009.07.033.

[56] J.S. Cristóvão, C.M. Gomes, S100 Proteins in Alzheimer’s Disease, Front. Neurosci. 13 (2019) 463. doi:10.3389/fnins.2019.00463.

[57] M.T. Heneka, M.P. Kummer, A. Stutz, A. Delekate, S. Schwartz, A. Vieira-Saecker, A. Griep, D. Axt, A. Remus, T.C. Tzeng, E. Gelpi, A. Halle, M. Korte, E. Latz, D.T. Golenbock, NLRP3 is activated in Alzheimer’s disease and contributes to pathology in APP/PS1 mice, Nature. 493 (2013) 674–678. doi:10.1038/nature11729.

[58] J. Flores, A. Noël, B. Foveau, J. Lynham, C. Lecrux, A.C. LeBlanc, Caspase-1 inhibition alleviates cognitive impairment and neuropathology in an Alzheimer’s disease mouse model, Nat. Commun. 9 (2018) 1–14. doi:10.1038/s41467-018-06449-x.

[59] C. Venegas, M.T. Heneka, Inflammasome-mediated innate immunity in Alzheimer’s disease, FASEB J. 33 (2019) 13075–13084. doi:10.1096/fj.201900439.

[60] H.E.R. Baughman, A.F. Clouser, R.E. Klevit, A. Nath, HspB1 and Hsc70 chaperones engage distinct tau species and have different inhibitory effects on amyloid formation, J. Biol. Chem. 293 (2018) 2687–2700. doi:10.1074/jbc.M117.803411.

[61] R.P. Koldamova, I.M. Lefterov, M.I. Lefterova, J.S. Lazo, Apolipoprotein A-I directly interacts with amyloid precursor protein and inhibits Aβ aggregation and toxicity, Biochemistry. 40 (2001) 3553–3560. doi:10.1021/bi002186k.

[62] A.R. Koudinov, T.T. Berezov, A. Kumar, N. V. Koudinova, Alzheimer’s amyloid β interaction with normal human plasma high density lipoprotein: association with apolipoprotein and lipids, Clin. Chim. Acta. 270 (1998) 75–84. doi:10.1016/S0009-8981(97)00207-6.

[63] A.M. Sandbichler, B. Jansen, B.A. Peer, M. Paulitsch, B. Pelster, M. Egg, Metabolic Plasticity Enables Circadian Adaptation to Acute Hypoxia in Zebrafish Cells, Cell. Physiol. Biochem. 46 (2018) 1159–1174. doi:10.1159/000489058.

[64] Y. Wu, D. Tang, N. Liu, W. Xiong, H. Huang, Y. Li, Z. Ma, H. Zhao, P. Chen, X. Qi, E.E. Zhang, Reciprocal Regulation between the Circadian Clock and Hypoxia Signaling at the Genome Level in Mammals, Cell Metab. 25 (2017) 73–85. doi:10.1016/j.cmet.2016.09.009.

[65] M. Egg, L. Köblitz, J. Hirayama, T. Schwerte, C. Folterbauer, A. Kurz, B. Fiechtner, M. Möst, W. Salvenmoser, P. Sassone-Corsi, B. Pelster, Linking Oxygen to Time: The Bidirectional Interaction Between the Hypoxic Signaling Pathway and the Circadian Clock, Chronobiol. Int. 30 (2013) 510–529. doi:10.3109/07420528.2012.754447.

[66] F. Bellanti, G. Iannelli, M. Blonda, R. Tamborra, R. Villani, A. Romano, S. Calcagnini, G. Mazzoccoli, M. Vinciguerra, S. Gaetani, A.M. Giudetti, G. Vendemiale, T. Cassano, G. Serviddio, Alterations of Clock Gene RNA Expression in Brain Regions of a Triple Transgenic Model of Alzheimer’s Disease, J. Alzheimer’s Dis. 59 (2017) 615–631. doi:10.3233/JAD-160942.

[67] D.P. Bartel, MicroRNAs: Target Recognition and Regulatory Functions, Cell. 136 (2009) 215–233. doi:10.1016/j.cell.2009.01.002.

[68] P. Kolasinska-Zwierz, T. Down, I. Latorre, T. Liu, X.S. Liu, J. Ahringer, Differential chromatin marking of introns and expressed exons by H3K36me3, Nat. Genet. 41 (2009) 376–381. doi:10.1038/ng.322.

[69] A. Dobin, C.A. Davis, F. Schlesinger, J. Drenkow, C. Zaleski, S. Jha, P. Batut, M. Chaisson, T.R. Gingeras, STAR: ultrafast universal RNA-seq aligner, Bioinformatics. 29 (2013) 15–21. doi:10.1093/bioinformatics/bts635.

[70] M. Pertea, D. Kim, G.M. Pertea, J.T. Leek, S.L. Salzberg, Transcript-level expression analysis of RNA-seq experiments with HISAT, StringTie and Ballgown, Nat. Protoc. 11 (2016) 1650–1667. doi:10.1038/nprot.2016.095.

[71] V. Wucher, F. Legeai, B. Hédan, G. Rizk, L. Lagoutte, T. Leeb, V. Jagannathan, E. Cadieu, A. David, H. Lohi, S. Cirera, M. Fredholm, N. Botherel, P.A.J. Leegwater, C. Le Béguec, H. Fieten, J. Johnson, J. Alföldi, C. André, K. Lindblad-Toh, C. Hitte, T. Derrien, FEELnc: A tool for long non-coding RNA annotation and its application to the dog transcriptome, Nucleic Acids Res. 45 (2017) gkw1306. doi:10.1093/nar/gkw1306.

[72] Y.J. Kang, D.C. Yang, L. Kong, M. Hou, Y.Q. Meng, L. Wei, G. Gao, CPC2: A fast and accurate coding potential calculator based on sequence intrinsic features, Nucleic Acids Res. 45 (2017) W12–W16. doi:10.1093/nar/gkx428.

[73] J.-C. Guo, S.-S. Fang, Y. Wu, J.-H. Zhang, Y. Chen, J. Liu, B. Wu, J.-R. Wu, E.-M. Li, L.-Y. Xu, L. Sun, Y. Zhao, CNIT: a fast and accurate web tool for identifying protein-coding and long non-coding transcripts based on intrinsic sequence composition, Nucleic Acids Res. 47 (2019) W516–W522. doi:10.1093/nar/gkz400.

[74] S.X. Ge, D. Jung, R. Yao, ShinyGO: a graphical gene-set enrichment tool for animals and plants, Bioinformatics. 36 (2019) 2628–2629. doi:10.1093/bioinformatics/btz931.

[75] Y. Liao, G.K. Smyth, W. Shi, featureCounts: an efficient general purpose program for assigning sequence reads to genomic features, Bioinformatics. 30 (2013) 923–930. doi:10.1093/bioinformatics/btt656.

[76] S.X. Ge, E.W. Son, R. Yao, iDEP: an integrated web application for differential expression and pathway analysis of RNA-Seq data, BMC Bioinformatics. 19 (2018) 534. doi:10.1186/s12859-018-2486-6.

[77] P. Langfelder, S. Horvath, WGCNA: an R package for weighted correlation network analysis, BMC Bioinformatics. 9 (2008) 559. doi:10.1186/1471-2105-9-559.

[78] P. Shannon, A. Markiel, O. Ozier, N.S. Baliga, J.T. Wang, D. Ramage, N. Amin, B. Schwikowski, T. Ideker, Cytoscape: a software environment for integrated models of biomolecular interaction networks., Genome Res. 13 (2003) 2498–504. doi:10.1101/gr.1239303.

[79] C.-H. Chin, S.-H. Chen, H.-H. Wu, C.-W. Ho, M.-T. Ko, C.-Y. Lin, cytoHubba: identifying hub objects and sub-networks from complex interactome, BMC Syst. Biol. 8 (2014) S11. doi:10.1186/1752-0509-8-S4-S11.

[80] B. Banerjee, D. Koner, R. Hasan, N. Saha, Molecular characterization and ornithine-urea cycle genes expression in air-breathing magur catfish (Clarias magur) during exposure to high external ammonia, Genomics. 112 (2020) 2247–2260. doi:10.1016/j.ygeno.2019.12.021.

[81] K.J. Livak, T.D. Schmittgen, Analysis of relative gene expression data using real-time quantitative PCR and the 2-ΔΔCT method, Methods. 25 (2001) 402–408. doi:http://dx.doi.org/10.1006/meth.2001.1262.

