## Supplementary material for "Genome-wide identification of novel long non-coding RNAs and their possible roles in hypoxic zebrafish brain": Suppl figure 1

### Slide 1
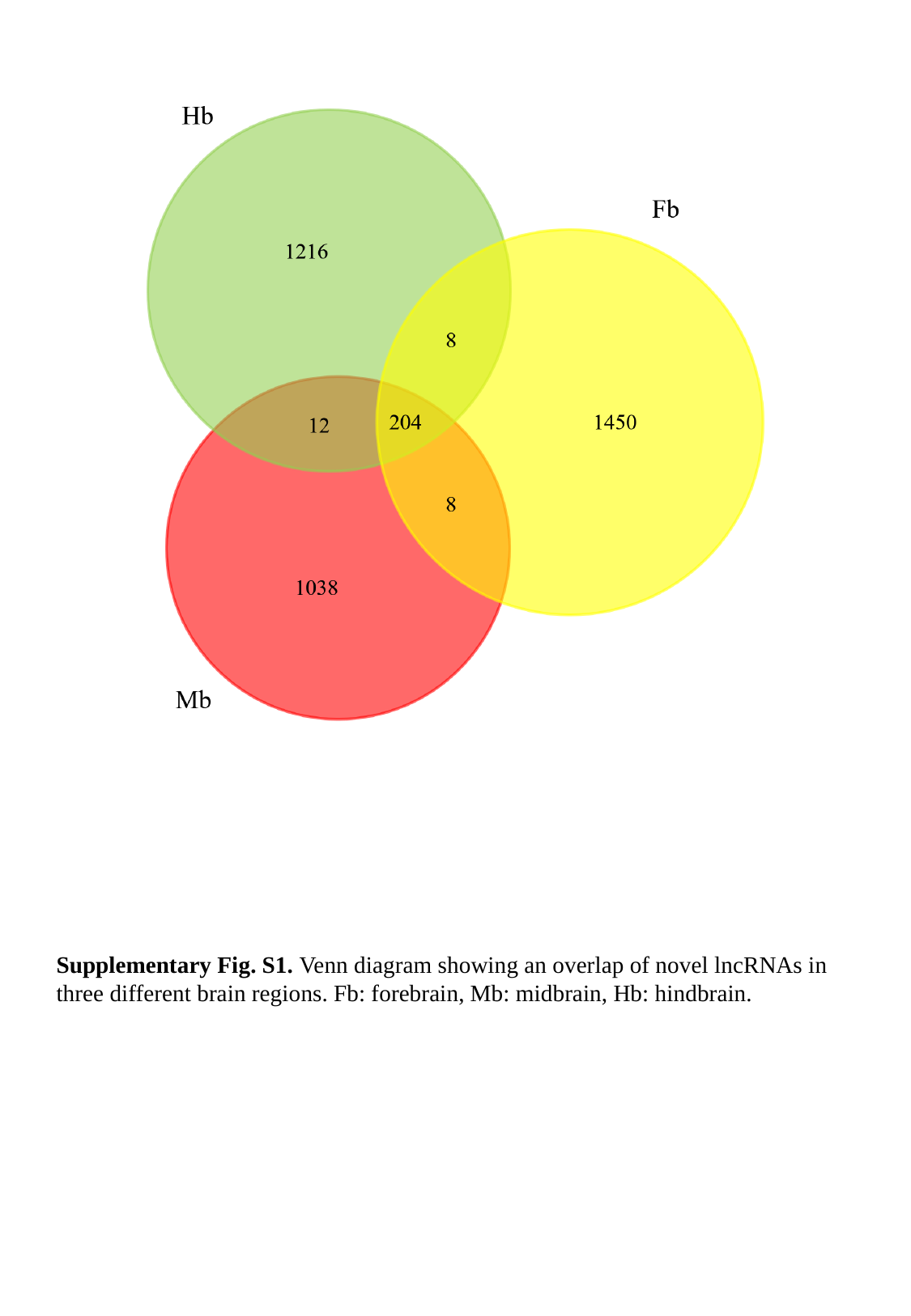

Supplementary Fig. S1. Venn diagram showing an overlap of novel lncRNAs in three different brain regions. Fb: forebrain, Mb: midbrain, Hb: hindbrain.

### Slide 2
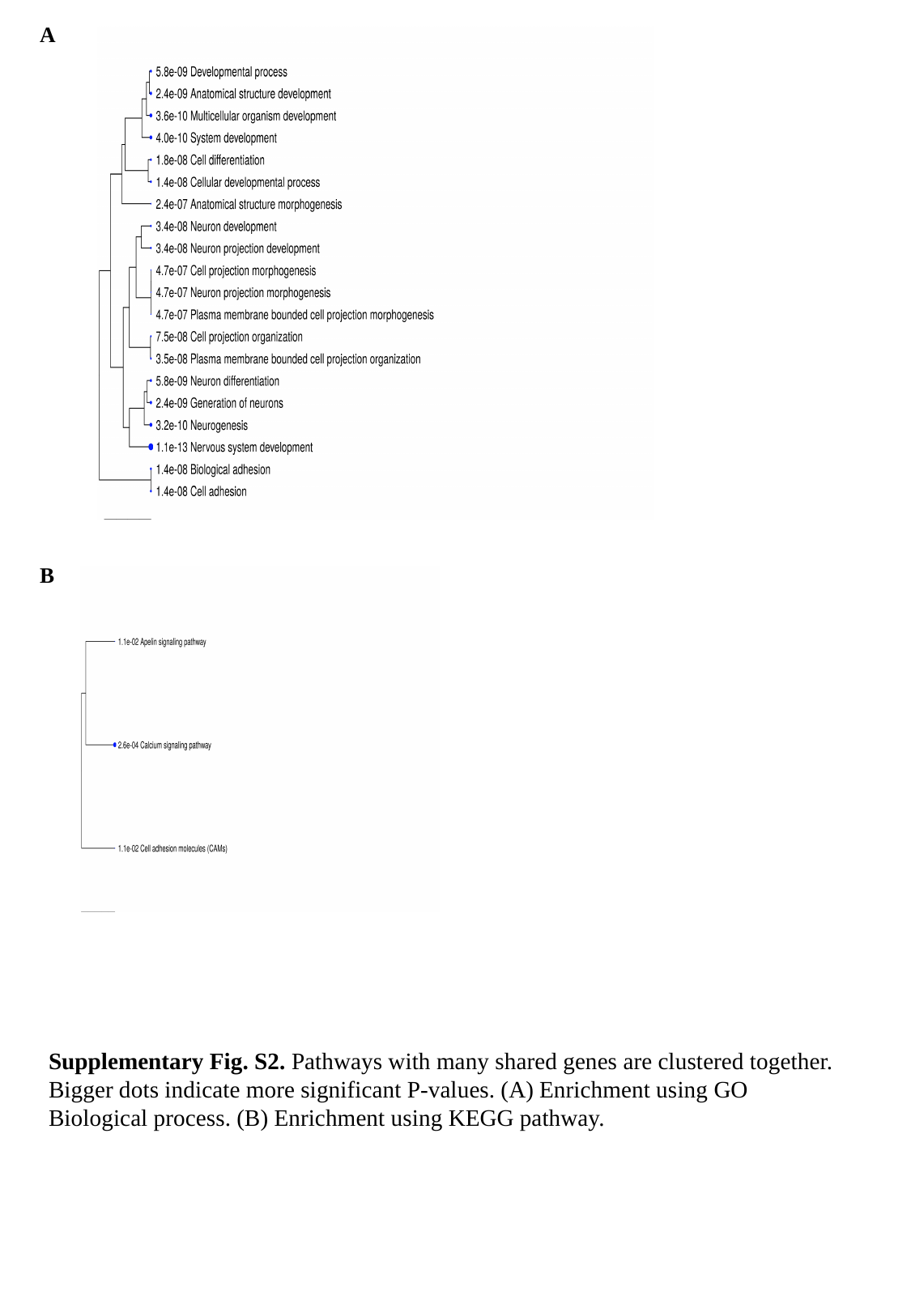

A
B
Supplementary Fig. S2. Pathways with many shared genes are clustered together. Bigger dots indicate more significant P-values. (A) Enrichment using GO Biological process. (B) Enrichment using KEGG pathway.

### Slide 3
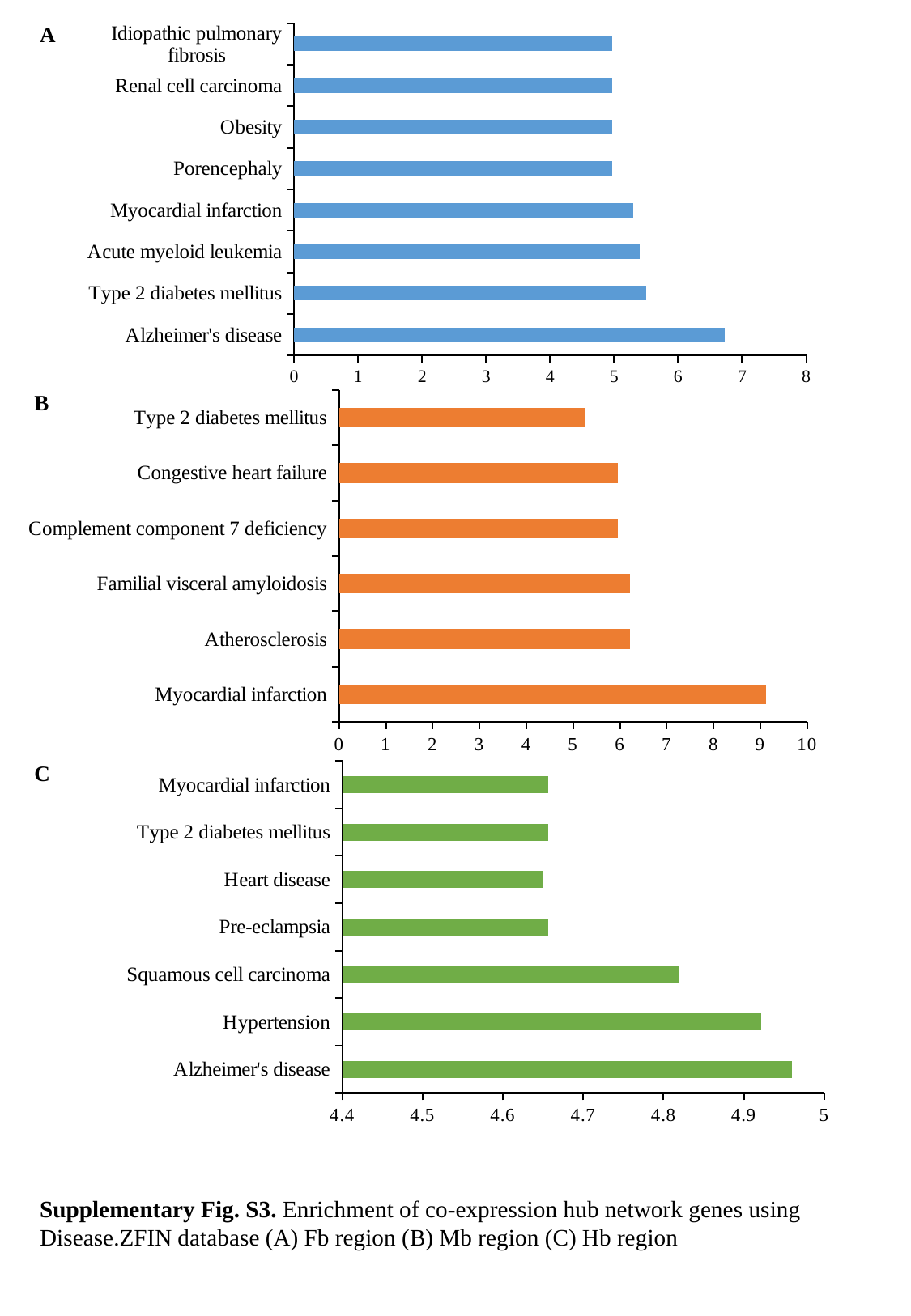

A
#### Chart
| Category | adj.Pval |
|---|---|
| Alzheimer's disease | 6.73 |
| Type 2 diabetes mellitus | 5.5 |
| Acute myeloid leukemia | 5.4 |
| Myocardial infarction | 5.3 |
| Porencephaly | 4.976233867378923 |
| Obesity | 4.976233867378923 |
| Renal cell carcinoma | 4.976233867378923 |
| Idiopathic pulmonary fibrosis | 4.976233867378923 |
#### Chart
| Category | adj.Pval |
|---|---|
| Myocardial infarction | 9.115030192171858 |
| Atherosclerosis | 6.215 |
| Familial visceral amyloidosis | 6.214 |
| Complement component 7 deficiency | 5.952243833954701 |
| Congestive heart failure | 5.9522438339547 |
| Type 2 diabetes mellitus | 5.259096653394756 |B
C
#### Chart
| Category | adj.Pval |
|---|---|
| Alzheimer's disease | 4.96 |
| Hypertension | 4.92198809308277 |
| Squamous cell carcinoma | 4.82 |
| Pre-eclampsia | 4.656463480375642 |
| Heart disease | 4.65 |
| Type 2 diabetes mellitus | 4.656463480375642 |
| Myocardial infarction | 4.656463480375642 |Supplementary Fig. S3. Enrichment of co-expression hub network genes using Disease.ZFIN database (A) Fb region (B) Mb region (C) Hb region
